# Spatial ligand–receptor gene annotations carry conditional heritability association across human tissues

**DOI:** 10.64898/2026.03.29.715138

**Authors:** Chen Yang, Xianyang Zhang, Jun Chen

## Abstract

Methods that map genetic risk to cells do not directly test whether spatially organized ligand–receptor (LR) gene annotations carry conditional heritability association. Here we introduce EdgeMap, which scores each gene by the spatial activity of its LR contexts in spatial transcriptomics data, maps these scores alongside cell-intrinsic annotations to SNP-level LD scores, and tests both jointly against GWAS summary statistics. Across 17 traits and five human tissues, edge ***z***-scores are systematically higher in biologically matched tissues: all 15 traits with both matched and unmatched tissue classes show this ordering (median **Δ*z* = 1.51**; sign ***P* = 3.1 × 10^−5^**), and 11 of 13 nominal-positive associations concentrate in the biologically matched set (***P* = 1 × 10^−4^**). The pattern survives broad LR-gene-class and cell-type composition controls. Donor-level cross-section summaries, independent GWAS replication for LDL and CAD, and cell-segmented Visium HD liver data provide convergent support. A secondary analysis prioritizes constituent genes within active LR contexts; sixteen of 31 prioritized genes are absent from standard gene-level methods. These results establish spatially weighted LR-gene annotations as a complementary layer for interpreting complex-trait genetic architecture.

## 1 Introduction

Genome-wide association studies (GWAS) have identified thousands of loci for complex traits, but linking these associations to the cellular programs through which they act remains a central challenge[1]. A major advance has come from partitioning SNP heritability across biologically informed annotations. Stratified LD score regression (S-LDSC)[2], which regresses GWAS association statistics on annotation LD scores, localizes heritability to functional categories; cell-type-specific extensions map heritability to trait-relevant tissues and cell populations[3, 4]; scDRS[5] resolves signal at single-cell resolution; and gsMap[6] extends this logic to spatial domains within tissues. These advances have sharpened heritability localization from tissues to cell types to spatial domains, but each still scores genes through cell-intrinsic activity rather than through the spatial interactions their products mediate between neighboring cells.

Yet tissues are not organized solely by cell identity; they are also shaped by communication between neighboring cells. More than 4,000 curated ligand–receptor (LR) interactions have been catalogued[7–9], and spatial transcriptomics now enables expression-based inference *in situ*[10–12]. A genetic effect need not remain within the cell in which a risk gene is expressed: a variant that alters a ligand, receptor or extracellular regulator can change the interface through which neighboring cells exchange signals. This biology is directly relevant to human genetics: interaction-network models of breast-cancer GWAS loci have found enrichment for interacting genes, including ligand–receptor interactions[13]; MR-CCC uses cis-eQTL instruments to infer causal cell–cell communication effects[14]; and Spacelink links spatially variable gene programs to disease genetics[15]. These studies provide convergent evidence that ligand–receptor gene organization shapes complex-trait risk.

Despite this convergent evidence, no existing framework constructs neighbor-aware, spatial LR-gene annotations for joint heritability analysis. S-LDSC and scDRS localize heritability using cell- or annotation-level scores; gsMap maps GWAS signal to genes through LD and proximity but does not aggregate LR expression scores across spatial neighbors. Communication inference tools such as CellChat[16] and CellPhoneDB[17] produce cell-pair activity scores suited for communication biology, but their outputs are not genome-wide SNP-level annotations. The conversion is not trivial: the aggregation rule determines which aspect of communication is testable in a heritability framework. This leaves a basic question unanswered: do spatially organized LR-gene annotations carry conditional heritability association beyond cell-intrinsic annotations?

Here we introduce EdgeMap, which tests **node** (cell-intrinsic) and **edge** (spatially weighted LR-context) annotations jointly (Fig. 1). From spatial transcriptomics data we derive two gene-level scores—cell-intrinsic expression and spatial LR-context activity—map both to SNP-level LD scores, and enter them jointly into S-LDSC, where each coefficient measures conditional association given the other annotations. A secondary analysis uses active LR contexts to prioritize their constituent genes. Because each context annotation is a scalar multiple of its membership vector, this analysis prioritizes genes within LR-labelled contexts but does not identify LR relations, direction or interaction effects (Methods). Across 17 complex traits and five human tissues, edge annotations carry systematically stronger conditional heritability association in biologically matched than unmatched tissues, a pattern that survives broad gene-class and composition controls; constituent-gene rankings within active LR contexts nominate candidates absent from standard gene-level methods.

**Fig. 1.**
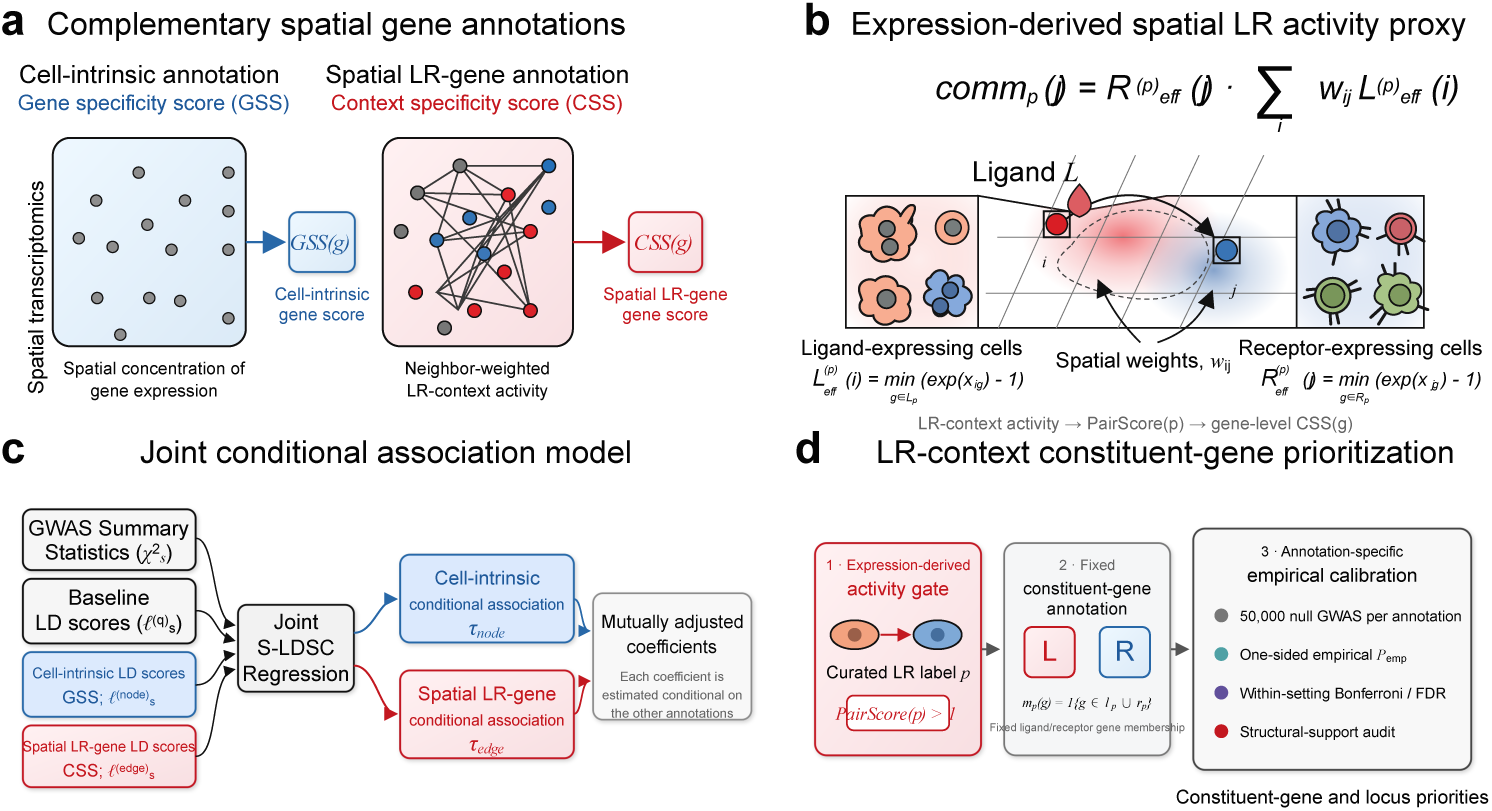
Overview of the EdgeMap framework. **a**, Spatial transcriptomics yields two complementary gene-level annotations: a cell-intrinsic gene specificity score (GSS) and a neighbor-weighted ligand–receptor context specificity score (CSS). **b**, Effective ligand expression in neighboring cells, spatial weights and receptor expression define an expression-derived LR-context activity proxy, which is summarized across locations and active contexts to obtain gene-level CSS. **c**, GSS- and CSS-derived LD scores enter joint S-LDSC with baseline annotations; the node and edge coefficients measure mutually adjusted conditional associations. **d**, In the secondary context-level analysis, expression-derived activity gates eligible LR labels, while fixed constituent-gene annotations are evaluated against annotation-specific empirical nulls and prioritized using within-setting calibration and structural-support auditing.

## 2 Results

### 2.1 The EdgeMap framework

EdgeMap takes spatial transcriptomics data and GWAS summary statistics as input (Fig. 1). For each tissue, we construct a spatial neighbor graph and compute an expression-derived LR score for each ligand–receptor (LR) label in the LIANA Consensus resource[7] (4,624 curated interactions; Methods). Existing communication inference tools[16, 17] estimate interaction activity at the level of cell pairs or individual channels—a format suited for communication biology but not for genome-wide heritability annotation. EdgeMap bridges this gap by deriving two parallel gene-level scores: the **gene specificity score** (GSS) quantifies how spatially concentrated a gene’s expression is; the **context specificity score** (CSS) summarizes an expression-derived, neighbor-weighted LR score; it is not a direct observation of signaling. GSS generates the node annotation and CSS the edge annotation; the two are entered jointly into the EdgeMap regression alongside baseline annotations. A positive edge coefficient *τ*_edge_ indicates a conditional association between SNPs near genes with high spatial LR scores and heritability given the node and baseline annotations. At this aggregate layer, EdgeMap asks whether genes with high spatial LR scores carry excess heritability as a class. Active LR labels provide a second layer: sparse constituent-gene annotations prioritize genes and loci, with empirical-null calibration and structural-support auditing to characterize the sparse-annotation regime (Methods). All context-level biological interpretation uses extracellular-verified pairs only (tiers 1 and 2; Methods, §4.35); full-database results are a sensitivity analysis (Supplementary Table 4).

### 2.2 Simulations support calibration and node–edge separation

We evaluated EdgeMap using real annotation LD scores from spatial transcriptomics with synthetic GWAS *χ*^2^ statistics across four architectures (null, node-only, edge-only, mixed; 500 replicates each; Fig. 2). Simulations were conducted on three tissues (heart, brain, gut) to test whether calibration generalizes across annotation structures (Supplementary Table 6, sheet “Multi-tissue Simulation”); results below are from heart.

**Fig. 2.**
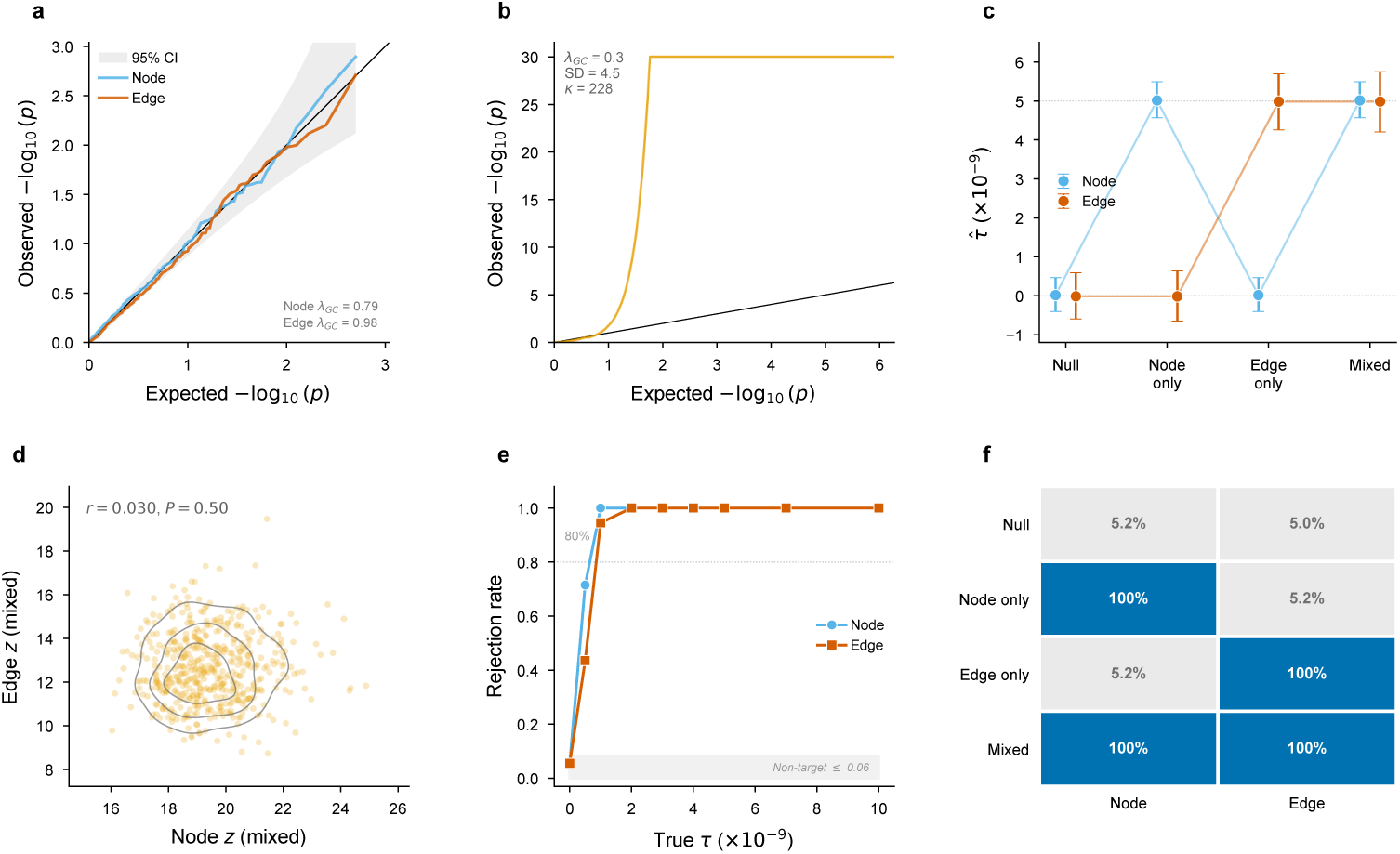
Simulation validation (heart annotation LD scores; brain and gut in Supplementary Table 6, sheet “Multi-tissue Simulation”). **a**, Aggregate null QQ plot (*n* = 500 replicates): node and edge *P* -values are well calibrated. **b**, Context-annotation null QQ plot: sparse constituent-gene annotations generate a heavy-tailed null that departs strongly from *N* (0, 1), motivating empirical calibration. **c**, Coefficient recovery: estimated *τ*^ (median *±* 95% interval) versus true *τ* across all scenarios; bias < 0.6%. **d**, Node and edge *z*-scores are uncorrelated under the mixed architecture (*r* = 0.030, *P* = 0.50), supporting separable estimation of the two components in simulation. **e**, Power curve: rejection rate as a function of true effect size *τ* ; the target component reaches 100% power by *τ* = 2 × 10^−9^ while the non-target component maintains nominal error. This idealized boundary excludes residual within-block LD; real-data detection limits are summarized from the observed standard errors in the text and Supplementary Table 2. **f**, Rejection-rate summary across null, node-only, edge-only, and mixed architectures, showing calibrated type-I error and complete power in the targeted non-null settings.

Under the null, type-I error was well calibrated: 5.2% for node and 5.0% for edge at *α* = 0.05 (*P* -values uniform by Kolmogorov–Smirnov test). Under node-only enrichment, the node component reached 100% power (mean *z* = 19.5) while the edge component maintained nominal error (5.2%). Under edge-only enrichment, the pattern reversed exactly: edge at 100% power (mean *z* = 13.6), node calibrated (5.2%). This separation supports node–edge separability in the tested simulation settings and the use of joint regression to distinguish conditional node- and edge-annotation signals. Null type-I error ranged from 4.4% to 6.2% across all three tissues (Supplementary Table 6, sheet “Multi-tissue Simulation”), indicating robust aggregate calibration across differences in annotation structure. An LD-aware null simulation— in which SNP-level noise terms are drawn from the empirical LD correlation matrix rather than independently—supports calibration under realistic LD structure at both aggregate and context-annotation levels (Methods; Extended Data Fig. 7g). As a direct calibration check, we conducted a genotype-derived null calibration battery using individual-level genotypes from the 1000 Genomes EUR Phase 3 panel (503 individuals). Under *τ*_edge_ = 0 across 22 configurations—spanning sparse causal (*π* = 0.001–0.1), infinitesimal, and node-enriched (5× enrichment) architectures, three trait-specific heritability and sample-size settings, and four heavy-tailed effect-size distributions (*t*_3_, *t*_5_, Laplace, point-normal)—context-annotation type-I error was 4.2– 5.3% at *α* = 0.05 across all architectures with uniform or node-only enrichment (1,000 replicates each; Extended Data Fig. 7d; Supplementary Table 6, sheet “Genotype Null Battery”). Under node-enriched architecture, the node component reached 78–82% power while the edge component maintained nominal error (4.8–5.1%), supporting node–edge separability with real genotype data. Heavy-tailed effects—including the *t*_3_ distribution—produced well-calibrated context-level error (4.3–5.1%). Under an LR-membership-enriched architecture (all ligand–receptor gene SNPs carry 5× enrichment but no context-specific enrichment), rejection was elevated to 8.1–8.3% in this deliberate confounding stress test, as expected from overlap between broad LR-class and sparse constituent-gene annotations. We therefore use LR-membership controls as a confounding sensitivity analysis. At the aggregate level, adding an LR-membership baseline retains 12 of 13 combinations; a membership-controlled empirical null provides a targeted sensitivity analysis for calibrated context-level rows (Supplementary Table 7; Extended Data Fig. 7c). LR-context annotations are much sparser (~1,000 nonzero SNPs per context), producing null *z*-scores that depart from *N* (0, 1) (sd ≈ 4.4, excess kurtosis 50–5,000; Fig. 2b). We therefore calibrate tail probabilities via 50,000-replicate empirical null simulation per setting (Methods, §4.8). Formal calibration was completed for 18 of 85 tissue–trait settings. This targeted subset contains all 12 aggregate nominal-positive settings in the four tissues for which these nulls were generated, plus six case-study or within-tissue comparators (brain AD and depression, gut CAD, and liver DBP, SBP and UC); hippocampal nulls were not generated. The subset was neither exhaustive nor selected to estimate a cross-setting discovery rate. Remaining context-level results are rankings only, and we do not report a study-wide discovery count. Right-tail empirical *P* -values preserve the signed *z*-score ordering within each calibrated context (mean Spearman *ρ* = 0.96 between signed-*z* rank and empirical-*P* rank across 18 empirical-null contexts; Extended Data Fig. 4c). Because these null *z*-distributions are shaped by annotation sparsity and are generally not centered at zero, the empirical *P* -value measures enrichment relative to that annotation’s own null rather than against *N* (0, 1), and the raw *z*-score is not by itself interpretable. The test is one-sided in the enrichment direction throughout (Methods, §4.8). We also audit structural support for each prioritized row by counting which ligand and receptor genes contribute autosomal SNP support and how many jackknife blocks carry the resulting annotation (Methods, §4.9; Supplementary Table 9). This check identifies which constituent genes and LD blocks carry each annotation.

To characterise detection limits, we performed a factorial power simulation across 190 parameter combinations (Extended Data Fig. 8). The simulation omits the residual correlation that LD induces among SNPs sharing a block, so the standard error it implies (3.4 × 10^−10^) is 4.9-fold smaller than the median obtained across the 85 real combinations (1.7 × 10^−9^; interquartile range 0.9–3.4 × 10^−9^; Supplementary Table 2).

We therefore state the detection boundary from the standard errors the analysis produced rather than from the simulation. Averaging power over the observed standard errors, 80% power is reached at *τ*_edge_ ≈ 6.7 × 10^−9^; at *τ*_edge_ = 10^−9^ average power is 26%. Effect size rather than sample size or heritability is what limits detection: at the reference effect size power is complete from *N*_eff_ = 100k upward and across *h*^2^ = 0.05–0.50 (Extended Data Fig. 8a,b). Power is largely insensitive to spatial resolution over the tested range: subsampling real heart Visium from 4,247 to 250 spots yields similar power (Extended Data Fig. 8e), because the CSS-annotated gene set (*∼*260 genes) and SNP-level annotation quality are stable across spot counts. Overlaying the 10 real tissue–trait results for which a rebuilt estimate exists on the *τ*_edge_ *×N*_eff_ landscape shows that significance tracks the ratio of *τ*_edge_ to that combination’s own standard error rather than *τ*_edge_ alone: heart–TG carries the largest effect of the ten (*τ*_edge_ = 1.1 × 10^−8^) and does not reach significance, because its standard error is correspondingly large (Extended Data Fig. 8a; Supplementary Table 2, sheets “Standard Errors” and “Detection Boundary”). This heterogeneity explains why the 13 of 85 threshold count is not an estimate of the prevalence of spatial LR-gene associations.

### 2.3 Spatial LR-gene annotation associations across traits and tissues

To characterize the aggregate edge-annotation landscape, we applied EdgeMap to 17 complex traits spanning cardiovascular, psychiatric, metabolic, and immune categories across five human tissues: heart (Visium, 4,247 spots, 479 active LR pairs), brain DLPFC (section 151507 from Maynard et al. 2021[18]; 4,226 spots, 346 active pairs), hippocampus (section V10B01-085 A1 from Thompson et al. 2025[19]; 4,992 spots, 691 active pairs), liver (section JBO001 from Guilliams et al. 2022[20]; 1,646 spots, 779 active pairs), and intestine (section B4; 2,575 spots, 231 active pairs; Fig. 3; Supplementary Table 2, sheet “Landscape”).

**Fig. 3.**
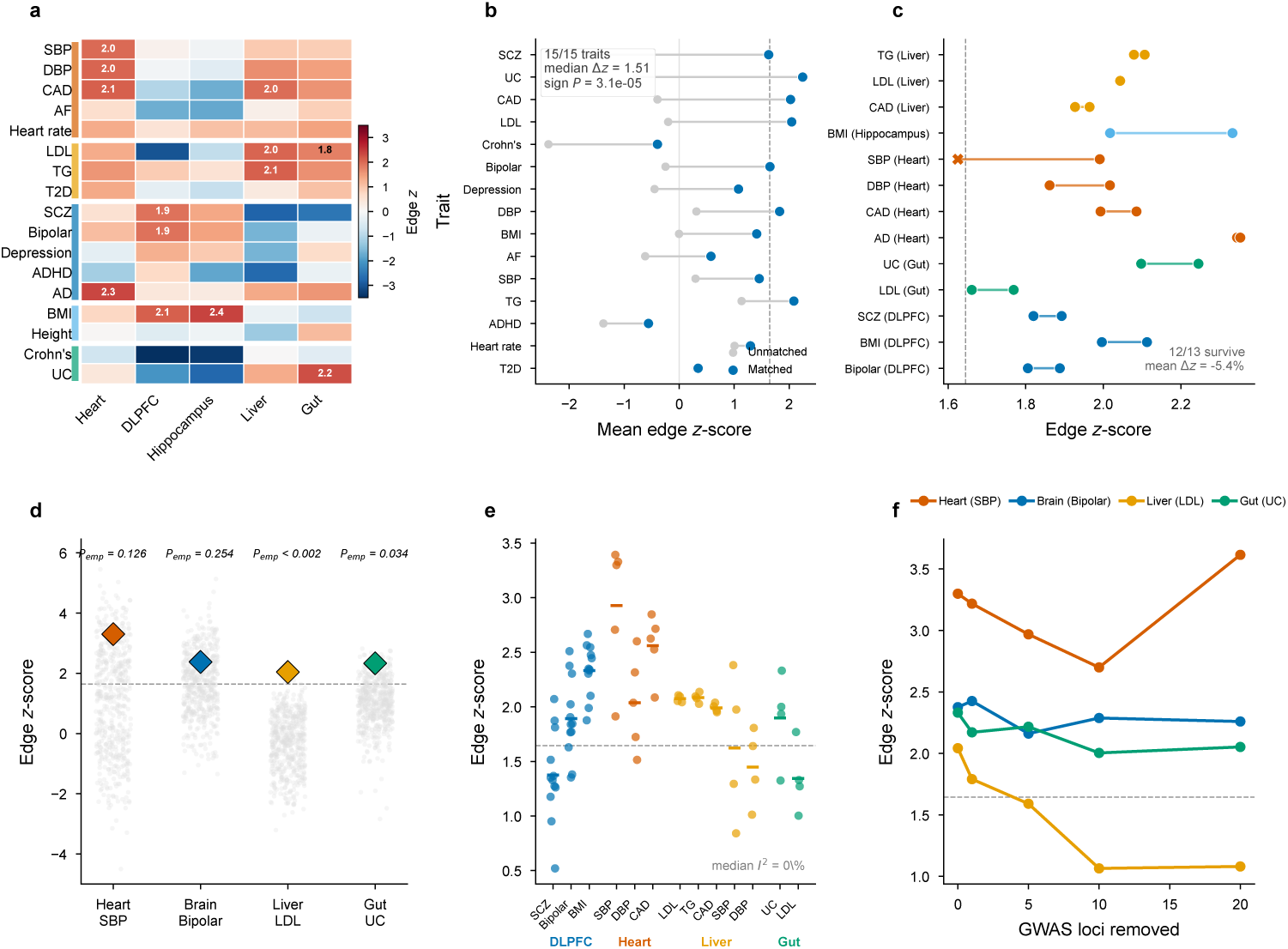
Aggregate spatial LR-gene annotation associations. **a**, Heatmap of edge *z*-scores across 85 trait–tissue combinations. Numbers mark nominal-positive combinations (unadjusted one-sided *P* < 0.05); none survives panel-wide FDR < 0.05. Colour reflects signed edge *z*-score. **b**, Within-trait biological coherence: for each trait, the mean edge *z*-score in biologically matched tissues is compared with the mean across unmatched tissues. All 15 traits with both classes show higher matched-tissue values (median Δ*z* = 1.51; one-sided sign *P* = 3.1 × 10^−5^). **c**, Cell-type composition control: paired edge *z*-scores before versus after conditioning on spatial-domain/composition annotations. Twelve of 13 tested nominal-positive combinations remain above the unadjusted one-sided *z* = 1.645 threshold after the additional control (mean Δ*z* = *−*5.4%). **d**, LR database specificity: real edge *z*-scores (diamonds) versus 500 expression-matched shuffled LR databases (dots). Liver (*P*emp < 0.002) and gut (*P*emp = 0.034) cross the unadjusted empirical *P* < 0.05 threshold; heart and brain do not. **e**, Cross-section robustness: edge *z*-scores across separate tissue sections (12 brain DLPFC sections from 3 donors, 5 heart sections from 5 donors, 4 liver sections from 4 donors, 4 gut sections from 4 donors). DLPFC serial sections are donor-correlated; the primary cross-section summary uses donor-level aggregation (Methods). **f**, Locus ablation: edge *z*-scores after removing the top *K* GWAS loci (*±*500 kb). Heart, brain, and gut remain above the nominal one-sided *z* = 1.645 threshold at all ablation levels; liver attenuates beyond 5 loci, consistent with concentrated lipoprotein-pathway architecture.

The clearest landscape-level signal appears before applying any significance threshold. For each trait with at least one biologically matched tissue and at least one unmatched tissue, we compared the mean edge *z*-score in matched tissues with the mean in the remaining tissues. The matched-tissue mean was higher for all 15 such traits (median Δ*z* = 1.51, mean Δ*z* = 1.58; one-sided sign *P* = 3.1 × 10^−5^; Supplementary Table 5). This paired within-trait contrast fixes GWAS power, polygenicity and the ligand–receptor gene universe; only the tissue annotation changes.

Thresholded results were consistent with the same pattern. Of 85 trait–tissue combinations, 13 crossed the unadjusted one-sided threshold (*P* < 0.05, one-sided)—3.1-fold over the ~4.25 expected by chance under a unit-normal reference. Combinations sharing a tissue or a trait are correlated, so the naive binomial *P* = 3.0 × 10^−4^ is too liberal; under a correlation model that accounts for shared tissue and trait structure (§4.10) the enrichment gives *P* = 2.7 × 10^−3^, and we quote that value throughout. Under the two-sided sensitivity analysis, 10 of these 13 retained a nominal-positive estimate, while 8 biologically unmatched combinations had nominal-negative edge estimates (negative *z*; e.g., Crohn’s disease in brain, *z* = −5.48; schizophrenia in liver, *z* = −2.80; Supplementary Table 5, sheet “Two-sided Inference”). Donor-level cross-section summaries (Extended Data Table 1; §2.5) supported aggregate spatial robustness. Quantile comparison of the aggregate *P* -values supports departure from the null for both annotations (Extended Data Fig. 6a).

Expressing the same results as standardised effect sizes rather than *z*-scores gives a measure comparable across annotations (§4.11). Meta-analysed across the 17 traits within each tissue, node *τ^∗^* is significant in all five tissues (*z* = 4.3–7.3) and edge *τ^∗^* in heart (*τ^∗^* = 0.13, *z* = 2.1, two-sided *P* = 0.033; Extended Data Fig. 6c,d). Averaging edge *τ^∗^* over all 17 traits mixes biologically matched and unmatched pairings; within matched pairings the values are substantial (liver–LDL 1.69, liver–TG 1.47, gut–UC 1.23, heart–CAD 1.05; Supplementary Table 2). We treat these values as within-study standardized summaries rather than as direct benchmarks against baseline-LD-conditioned S-LDSC annotations, because the present regression uses the gsMap quick-mode baseline. Individual trait–tissue *z*-scores are moderate (*z* ≈ 2–3), as expected when aggregate annotations carry limited per-SNP variance in the S-LDSC regime[2]; no combination survives Benjamini–Hochberg correction across the panel at FDR < 0.05 (*q*_min_ = 0.21). We refer to the 13 combinations with one-sided nominal *P* < 0.05 as the *nominal-positive* set; the aggregate claim rests on pattern-level tests of within-trait tissue coherence and matched-tissue concentration across the full panel, not on any single combination.

We next assessed whether this pattern was concentrated in biologically matched tissues. Biological matching is defined by a prespecified rule — cardiovascular traits in heart, psychiatric traits and BMI in brain and hippocampus, metabolic and cardiovascular traits in liver, inflammatory bowel disease in gut (Methods, §4.10) — which covers 24 of the 85 combinations. Of the 13 above-threshold combinations, 11 fall inside that matched set and two fall outside it (heart–AD and gut–LDL). Depletion shows the complementary trend but is not deterministic: most depleted combinations lie outside the matched set, while hippocampus–ADHD is a biologically matched pairing with negative edge *z*. We therefore interpret the pattern as a distributional concentration rather than as an all-or-none directional rule. The conditional controls below address whether this coherence reduces to generic gene importance or other tissue-correlated properties. The same contrast holds in an unpaired rank analysis across all 85 combinations: edge *z* is higher in biologically matched pairings than in the rest (Mann–Whitney *P* = 6.3 × 10^−6^; median *z* 1.52 against 0.05; Fig. 3b), a weighted *z*-score combination (Stouffer’s method) over the 24 gives *z* = 6.25, and the signal remained significant after removing any single tissue (leave-one-tissue-out *P* < 6 × 10^−4^ for all five). Trait-label permutation, which preserves the cross-trait dependence structure within each tissue, confirmed the concentration: shuffling labels 10,000 times, the null expectation was 3.8 nominal-positive threshold crossings in biologically matched pairings versus the 11 observed (*P* = 1 × 10^−4^; Δ*z* = 1.46, never matched under permutation). A narrower sensitivity enumeration strengthens every statistic (Stouffer *z* = 7.12, Mann–Whitney *P* = 3.0 × 10^−8^; Supplementary Table 5); we report the broader rule because it is implied by the tissue–trait categories defined above. The joint analysis also revealed heterogeneous node–edge profiles. Atrial fibrillation in heart had a strong node estimate (*z*_node_ = 6.99) but no aggregate edge enrichment (*z*_edge_ = 0.58); T2D in liver showed the same qualitative contrast (*z*_node_ = 2.20, *z*_edge_ = 0.34). Height, despite being a well-powered highly polygenic trait, was also edge-null across tissues (maximum *z* = 1.11). These contrasts indicate that the edge annotation does not increase uniformly with node signal or GWAS power. Three further controls bound the interpretation of these findings. Permuting the pairing structure of the database—retaining every active ligand and receptor but randomly reassigning which ligand is paired with which receptor—left aggregate edge *z*-scores unchanged (empirical *P* = 0.08–1.00 across nine tissue–trait combinations; §4.31). This is the expected behaviour of a statistic that scores each gene by its highest expression-derived LR-context score and so summarises the spatial organisation of the LR gene class as a whole. Thus the aggregate statistic is insensitive to LR pair labels. Holding each ligand fixed and resampling 500 alternative receptors, real pairings achieved systematically higher context specificity than random ones (mean empirical *P* = 0.435 against a null expectation of 0.5; *P* = 1.3 × 10^−28^), and the effect grew monotonically with context specificity, from 5.9% of pairs significant at specificity ≥ 2 to 22.2% at specificity ≥ 7. This partner-resampling diagnostic evaluates the specificity of the spatial score and the active-context gate. Replacing the LIANA Consensus database with 500 expression-matched shuffled databases placed the observed edge statistic beyond the unadjusted empirical threshold in liver and gut, but not in heart or brain (Fig. 3d; §4.29; Discussion). Re-running the full pipeline with four alternative LR databases from the LIANA resource (CellTalkDB, CellChatDB, CellPhoneDB and their core intersection; 42–442 mean active pairs) yielded concordant aggregate rankings (Spearman *ρ* = 0.36–0.97 versus Consensus). Sequential locus ablation supported polygenic distribution in heart, brain, and gut; liver attenuated beyond five loci, consistent with concentrated lipoprotein-pathway architecture (Fig. 3f; §4.34).

The signal is also not explained by broader gene classes that are themselves heritability-enriched. Adding annotations for all genes, for all expressed genes, for the full ligand–receptor database and for the most highly expressed decile—separately and all four at once— retained 12 of the 13 nominal-positive combinations. With all four included at once the mean edge *z*-score falls by 6.3%, as expected when annotations covering every gene absorb part of the same variance. The broad ligand–receptor geneclass annotation, the most directly overlapping gene-class control, retains 12 of 13 and raises the mean edge *z*-score by 5.3% (§4.30; Supplementary Table 5, sheet “Added Baselines”). The same table reports a stronger diagnostic conditioning on tissue-active LR genes; because that indicator shares active LR-gene support with the edge annotation, it is a heavily conditioned sensitivity analysis rather than a broad gene-class baseline.

### 2.4 Node and edge annotations show distinguishable conditional signals

A natural concern is whether the edge annotation merely re-identifies the same genes that drive node heritability. A more direct test is at the level of heritability rather than of gene lists: in simulations where heritability is concentrated entirely in one simulated annotation, the other returns a near-zero *z*-score in both directions (§2.2; Fig. 2d,f), so the joint regression distinguishes the two conditional signals even when the underlying gene sets are correlated. Gene-level composition is concordant with this. Comparing genes in prioritized LR contexts with the top-ranked genes by GSS (top 100 per tissue, e.g. TTN, MYH7 and ACTA1 in heart; Methods), across the trait–tissue combinations that contained calibrated context-level priorities, the mean Jaccard overlap was 0.002 (median 0), with only 2.2% of prioritized constituent genes appearing among the top node genes (Extended Data Fig. 3a). Prioritized constituent genes showed only weak node-score enrichment (mean rank-biserial correlation = 0.06; Mann–Whitney *P >* 0.05 in 12 of 13 combinations), and genome-wide GSS and CSS were essentially uncorrelated (mean Spearman *ρ* = 0.02). GSS is also distinct from generic spatial autocorrelation: across 15,217 heart genes it correlates negatively with Moran’s *I* (Spearman *ρ* = −0.12; Extended Data Fig. 7a). A coordinate-noise stress test gave the same separation: at one-neighbour displacements, GSS rankings collapse whereas CSS rankings remain stable (Extended Data Fig. 7b), consistent with the different supports of the node and edge scores rather than node-to-edge leakage. All 13 nominal-positive edge combinations also have nominal-positive node estimates. This co-occurrence motivates the explicit node–edge distinguishability analyses below. Together with the simulation results (§2.2), these analyses show that the two fitted annotations carry distinguishable conditional signals and produce largely non-overlapping gene priorities (Extended Data Fig. 3a).

This complementarity extends beyond EdgeMap’s own node component. Comparing prioritized constituent genes against gene-level association testing analogous to MAGMA[21] (minimum SNP *P* within ±10 kb, Bonferroni threshold *P* < 2.5 × 10^−6^) and Open Targets Locus-to-Gene fine-mapping[22] (L2G score ≥ 0.5, matched per trait; Methods), 16 of 31 unique prioritized genes (52%) were identified by neither method (Extended Data Fig. 5; Supplementary Table 4, sheet “MAGMA-L2G”).

These include genes with extracellular or disease-relevant biology not identified by either gene-level method: NECTIN1, an adhesion receptor in the highest-ranked CADM3–NECTIN1 context for bipolar disorder in brain; GPC3, a glypican coreceptor whose loss-of-function causes an overgrowth syndrome with congenital cardiac defects[23], identified through a GPC3–LRP1 receptor-axis candidate for coronary artery disease in heart; and F9 (coagulation factor IX), a hepatocyte-secreted coagulation factor whose activated form can interact with LRP1[24], identified here through a liver–LDL F9–LRP1 endocytic-clearance candidate (§2.7). None of these genes reaches the stated gene-level significance threshold in the comparison analysis. Their selection here reflects a different criterion: constituent genes of empirically prioritized LR-labelled annotations, rather than proximity to a lead variant alone.

We also tested whether the aggregate edge association was explained by spatial variation in cell-type composition. Augmenting the S-LDSC regression with spatial domain annotations that capture local cellular composition (published DLPFC cortical layers and expression-derived Leiden domains in the other tissues; Methods, §4.28) had minimal impact: 12 of the 13 nominal-positive combinations retained nominal significance. Heart–SBP remained positive but moved from *z* = 1.99 to *z* = 1.63, just below the one-sided threshold; the mean *z*-score change was 5.4% (Supplementary Table 5, sheet “Primary Domain Control”). The aggregate edge pattern is therefore not explained solely by these composition gradients.

### 2.5 Aggregate spatial LR-gene associations show cross-section robustness and independent-GWAS support

To test whether edge signals reflect stable tissue biology rather than section-specific artifacts, we ran EdgeMap independently on multiple sections per tissue (12 brain DLPFC from 3 donors[18], 5 heart from 5 donors[25], 4 liver from 4 donors[20], 4 gut from 4 donors; Fig. 3e). Eleven of the 13 nominal-positive aggregate combinations have more than one section and could be tested; heart–AD was not part of the replication panel and hippocampus–BMI has a single section, so no cross-section robustness claim is made for those two. All 11 showed donor-level aggregate support (Extended Data Table 1). Sections within a donor are not independent — the 12 DLPFC sections are four serial sections from each of three donors — so we combine at donor level rather than section level. Donor-level meta-analytic *z*-scores range from 2.25 (brain SCZ) to 5.72 (heart CAD), all m<u>eta</u> *P* < 0.013; the section-level combination used previously is larger by up to 12*/*3 = 2 for brain and is reported alongside it. Section-level edge *z*-scores were consistently positive across separate tissue sections (Extended Data Fig. 3c). Raw communication-score inputs showed moderate concordance across independent heart sections (mean Pearson *r* = 0.66 across the six unique Kuppe section pairs; Extended Data Fig. 4a), before entering S-LDSC. Context-level *z*-score vectors agree across sections at *ρ* = 0.98–0.99 (Supplementary Table 5, sheet “Scale Invariance Audit”); however, scalar invariance (§2.1) means this agreement reflects shared gene-set and LD structure rather than independent measurement. Cross-section support for the aggregate signal rests on donor-level meta-analysis and on the consistent sign of section-level estimates. A predefined cross-section robustness and replication-diagnostic protocol (Methods; Extended Data Fig. 7f) supported these findings. Aggregate edge *z*-scores were positive in all 71 section–trait observations across the 11 replicable combinations. Sections are not independent, so we quote the sign test at the level at which the units are: 44 of 44 donor–trait observations positive, *P* = 5.7 × 10^−14^, and 11 of 11 combination-level means positive, *P* = 4.9 × 10^−4^; the section-level figure (*P* = 4.2 × 10^−22^) is reported for completeness but assumes section independence. Sign concordance among the top 20 pairs was 100% across all 41 section pairs, and the top-20 pairs from any one section showed 17× enrichment for top-20 status in every other section; as with the context-level correlations above, scalar invariance means these statistics characterise the annotation rather than independent measurement.

#### Independent GWAS replication

Replacing the primary LDL GWAS (UK Biobank) with a fully independent dataset (GLGC 2013[26]; *N* = 173,082; no sample overlap), the liver edge signal replicated (*z* = 2.24, *P* = 0.013), with LR-context constituent-gene *z*-scores concordant between the two GWAS (Spearman *ρ* = 0.53, *P* = 1.7 × 10^−50^; Fig. 5d; Extended Data Fig. 4b) and PCSK9–SORT1 among the top-ranked contexts in both. An analogous test for heart using FinnGen Release 10 CAD[27] (*N*_eff_ = 166,436; no sample overlap) also replicated (*z* = 1.94, *P* = 0.026).

#### Cross-platform support

EdgeMap applied to cell-segmented Visium HD liver data[28] (8 *µ*m resolution; independent donors and laboratory) supported the edge signal (*z* = 1.93, *P* = 0.027; Fig. 5e). At native 8 *µ*m resolution, five highlighted liver–LDL maps showed distinct primary sender and receiver cell types—including hepatocyte F9 paired with stellate-cell LRP1, an endocytic-clearance candidate, and hepatocyte C3 paired with cholangiocyte C3AR1 (Fig. 5c)—an expression-derived spatial allocation consistent with intercellular rather than purely autocrine geometry.

### 2.6 Heart aggregate associations and cardiovascular gene contexts

We next examined LR-context constituent-gene results in individual tissues, using empirical null calibration where available (§2.2, §4.8). Thresholds and rankings are reported within each calibrated context and do not define a study-wide discovery set. Three cardiovascular traits in heart—SBP, DBP and CAD—cross the unadjusted threshold (SBP *z* = 1.99, DBP 2.03, CAD 2.09). Heart–AD also reaches threshold (*z* = 2.34), but is treated separately below as an out-of-rule AD signal. We examined the constituent-gene rankings within these nominal-positive combinations.

For systolic blood pressure (SBP, *N* = 340,159), the node annotation was highly significant (*z* = 9.46, *P* = 1.5 × 10^−21^), consistent with prior findings[6]. The edge annotation had a positive conditional estimate (*z* = 1.99, *P* = 0.023; *τ*_edge_ = 2.99 × 10^−9^, SE = 1.50 × 10^−9^).

Exploratory LR-context analysis of SBP (370 active contexts, 367 empirically testable; empirical null calibration, Methods) prioritized VEGFD–ITGA9 at the within-combination Bonferroni threshold (*P*_emp,Bonf_ = 0.007; Fig. 4a; Supplementary Table 3, sheet “Heart”), a VEGF-D–integrin *α*9*β*1 vascular/lymphangiogenic candidate[29]. Structural QC shows that this constituent-gene annotation is driven by autosomal support for ITGA9, because the ligand side does not contribute SNP support in the current autosomal weight matrix; we therefore use it as pathway-level prioritization rather than as evidence for a VEGFD–ITGA9 relation (Supplementary Table 9). Other high-ranked extracellular-verified contexts spanned complement, fibronectin–ECM, Notch vascular signalling (including JAG2/THBS2–NOTCH3; *NOTCH3* mutations cause CADASIL[30, 31]), GPCR–cAMP signalling, and lipoprotein receptor signalling. The high-ranked rows span several known cardiovascular gene families.

**Fig. 4.**
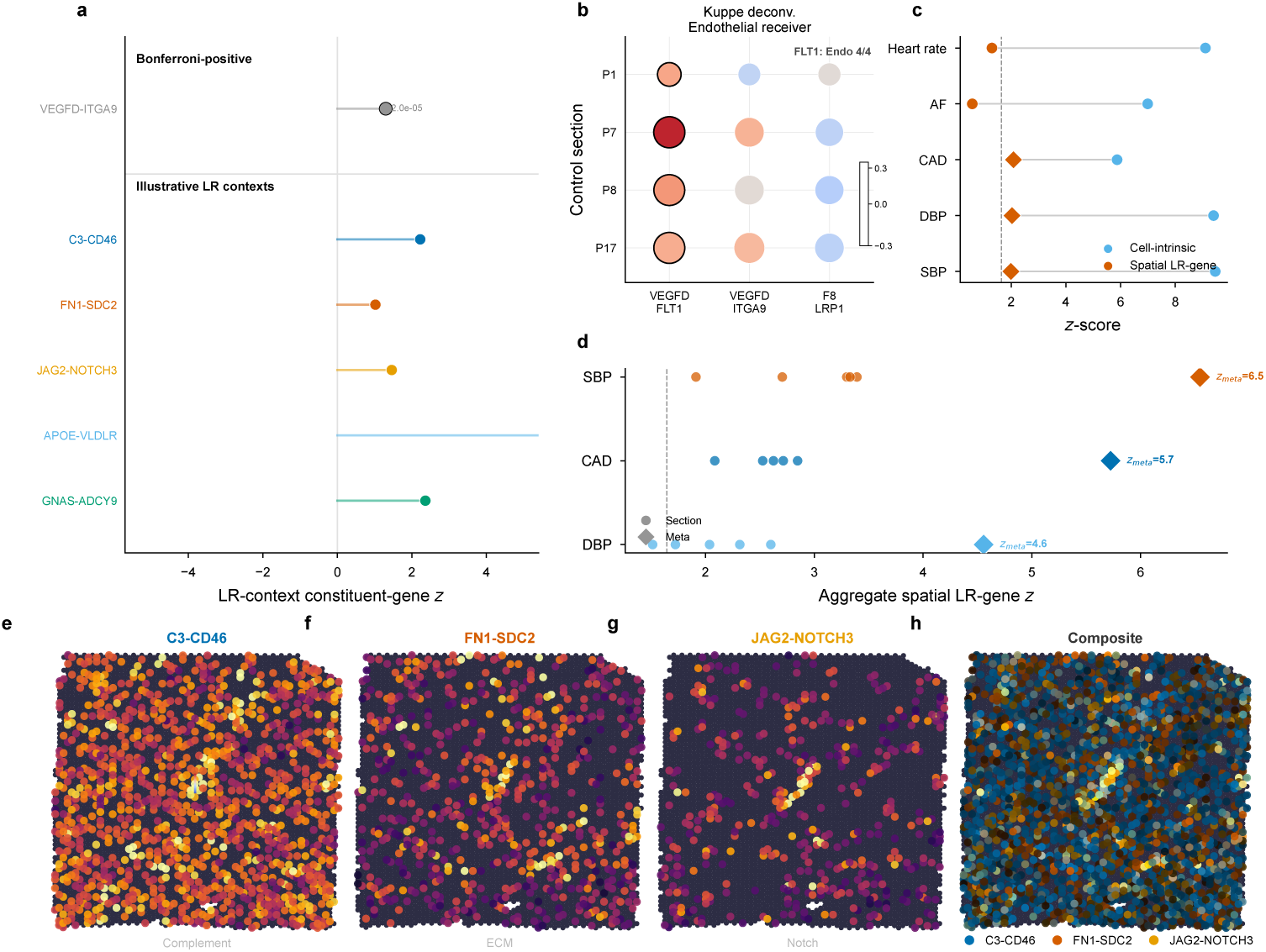
Heart aggregate conditional associations and LR-context constituent-gene summaries. **a**, Heart–SBP statistics for selected LR-context constituent-gene annotations. VEGFD– ITGA9 is the sole within-combination Bonferroni-positive row among 367 tested annotations; the lower band shows five prespecified illustrative contexts. Numbers beside Bonferroni-positive rows are one-sided empirical *P* values. **b**, Expression-based localization of three selected contexts across four Kuppe control-heart sections. Bubble size increases with the fraction of context activity assigned to endothelial receivers, colour denotes log_2_ enrichment over the cell-type composition expectation, and black outlines indicate that the receptor has its highest mean expression in the endothelial compartment. FLT1 meets the latter criterion in all four sections. **c**, Joint conditional S-LDSC *z* statistics for cell-intrinsic and spatial LR-gene annotations across five cardiovascular traits. Orange diamonds mark nominal-positive spatial LR-gene coefficients at one-sided *P* < 0.05; these *P* values are unadjusted across the 85 tissue–trait tests. **d**, Section-level aggregate spatial LR-gene statistics for SBP, CAD and DBP across five heart sections. Circles denote individual sections, diamonds denote the equal-weight Stouffer combination of the five section statistics, and the dashed line marks the nominal one-sided threshold *z* = 1.645. **e–g**, Expression-derived spatial LR-score maps for C3–CD46, FN1– SDC2 and JAG2–NOTCH3. Colour represents log_1_*p*-transformed activity scaled within each map. **h**, RGB overlay of the three independently normalized maps. Context-level statistics in panel **a** use fixed constituent-gene annotations (§2.1).

Diastolic blood pressure (*z* = 2.03) and coronary artery disease (*z* = 2.09) were also nominal-positive in heart. Their context rows again concentrated on vascular and endocytic receptor axes (VEGFD–ITGA9, the FLT1 receptor-side VEGFD–FLT1 row, F8–LRP1 and GPC3–LRP1), though structural QC flags several as single-sided (only one constituent gene contributing autosomal SNP support) or single-block annotations. The same receptor axes recur across all three cardiovascular traits. Atrial fibrillation and heart rate, by contrast, showed no edge signal in the same tissue (Fig. 4c). We then asked whether these spatial contexts localize to plausible cardiac compartments using the published Kuppe control-heart Visium deconvolution weights across four independent sections (P1, P7, P8 and P17)[25]. The CAD-leading VEGFD–FLT1 row showed the clearest vascular receiver signature: endothelial receiver compartments captured a median 23.7% of expression-derived LR activity and were enriched 1.12-fold over the cell-type composition expectation, and FLT1 mean expression was highest in the endothelial compartment in all four sections (Fig. 4b; Supplementary Table 8). Because VEGF-D is classically linked to VEGFR2/KDR and VEGFR3/FLT4 rather than VEGFR1/FLT1, we read this as FLT1 receptor-side vascular localization rather than evidence that VEGF-D directly binds FLT1. VEGFD–ITGA9 showed weaker but directionally consistent endothelial receiver support (median enrichment 1.05). F8– LRP1 localized differently: F8 was endothelial-enriched in all four sections, whereas LRP1 localized preferentially to fibroblast, myeloid or vascular smooth-muscle compartments, consistent with a coagulation–clearance interface rather than an autocrine endothelial signal. Expression-derived maps for C3–CD46, FN1–SDC2 and JAG2– NOTCH3 illustrate distinct regional activity patterns and their overlay (Fig. 4e–h); these maps visualize the spatial proxies and are not tests of the named LR relations. Alzheimer’s disease also crossed the unadjusted aggregate threshold in heart (*z* = 2.34, *P* = 0.010). The row with the smallest within-combination Bonferroni-adjusted value is HLA-A–ERBB2 (*P*_emp,Bonf_ = 0.007), but this row is structurally single-sided after MHC filtering and is not used as the mechanistic AD anchor. APP–LRP1 also appears in the FDR set (*P*_emp_ = 2.8 × 10^−4^, FDR = 0.051), with both constituent genes mappable across multiple LD blocks. This overlap is not treated as a mechanistic anchor or as evidence for an APP–LRP1 relation (Supplementary Table 9).

All three cardiovascular traits showed section-level support for the aggregate signal across five heart sections from separate donors (Fig. 4d; §2.5). The edge signal was robust across spatial neighborhood sizes *k ∈ {*4, 6, 10, 15, 20*}* and context-specificity thresholds *P*_%_ 85, 90, 95, 99 (nominal-positive in all 20 tested parameter combinations; Extended Data Fig. 3b). As discussed in §2.5, cross-section aggregate robustness is summarized through donor-level meta-analysis and consistent positive estimates, not through context-level rank concordance alone.

### 2.7 Liver aggregate associations and clearance-receptor gene contexts

Liver offers a different test: because LDL biology includes unusually well-characterized genetic and therapeutic mechanisms, including PCSK9 and the 1p13/SORT1 locus, liver–LDL provides a positive biological reference point for ranking LR-labelled constituent-gene candidates rather than a formal single-channel validation. LDL cholesterol showed a positive aggregate edge association (*z* = 2.04, *P* = 0.021). LR-context analysis with empirical null calibration identified four rows at the within-combination Bonferroni threshold among 664 tested (668 active; Fig. 5a; Supplementary Table 3, sheet “Liver”). Structural QC indicates that these rows are single-sided or single-block annotations (Supplementary Table 9). They therefore nominate endocytic and clearance-receptor loci—including LRP1, IGF2R and SDC1—rather than providing direct support for each ligand–receptor pairing. This interpretation is consistent with the pathway context: in mouse genetic studies, syndecan-1 is the primary hepatocyte proteoglycan receptor mediating triglyceride-rich lipoprotein clearance[32], and LDLR/LRP1-family endocytic receptors contribute to hepatic ligand and remnant-lipoprotein clearance. Within the broader context ranking, PCSK9–SORT1 had both constituent genes mappable and ranked among the leading LDL contexts (*P*_emp_ = 0.021; not individually Bonferroni-significant) within a well-characterized LDL regulatory pathway: SORT1 facilitates intracellular trafficking and secretion of PCSK9[33], evolocumab and alirocumab target PCSK9, and PCSK9 promotes LDLR degradation in hepatocytes. The SORT1 locus (1p13.3) is one of the strongest common-variant signals for LDL cholesterol[34]. Spatial maps illustrate this ranking: PCSK9–SORT1 and ANGPTL4–SDC4 show distinct expression-derived spatial LR-score patterns, while LRPAP1–LRP1 is included as a tier 3 contextual LRP1 clearance-receptor locus map reported for completeness rather than as part of the extracellular-verified biological interpretation (Fig. 5f–h). The composite overlay separates these spatial patterns directly (Fig. 5i).

**Fig. 5.**
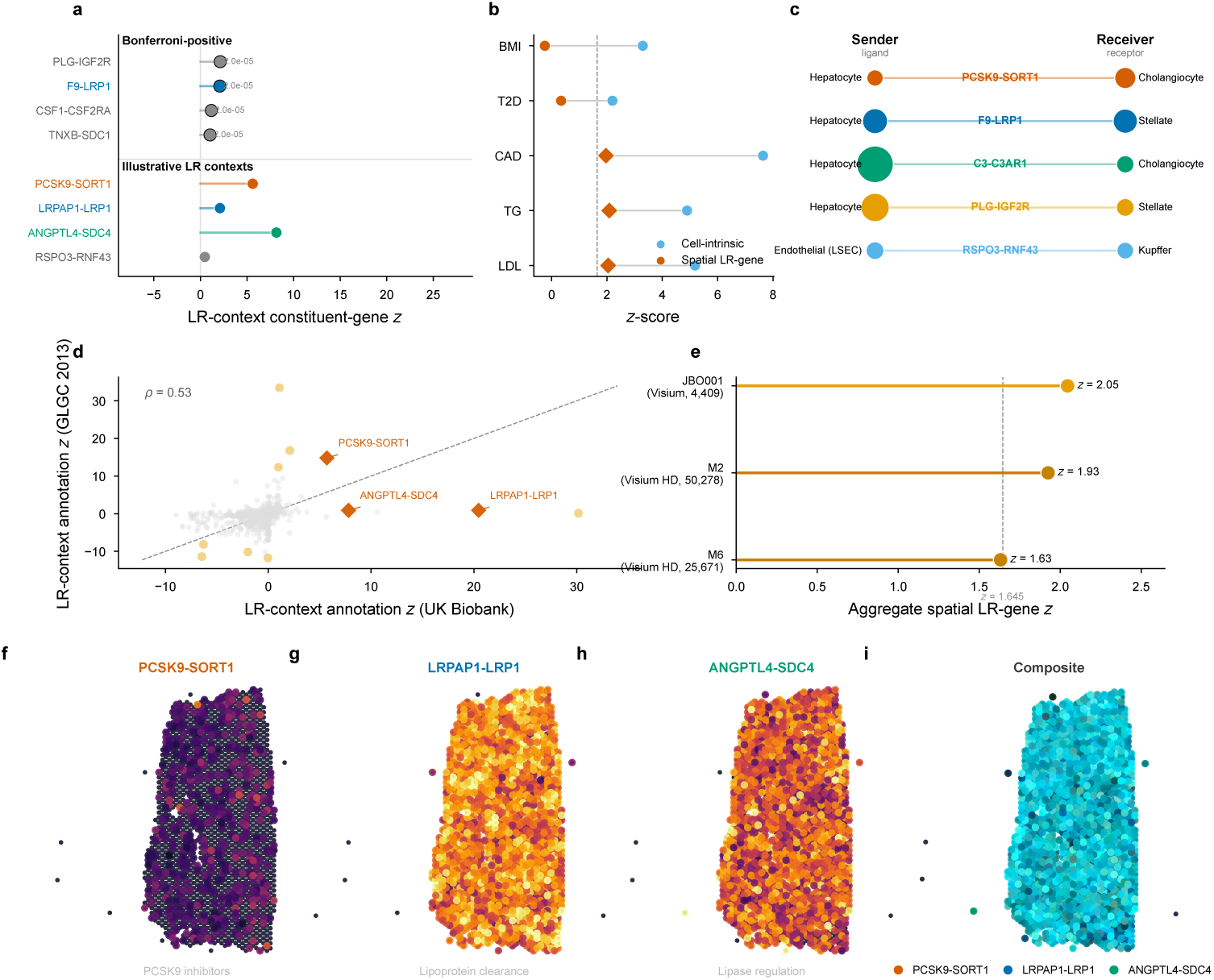
Liver aggregate conditional associations and LR-context constituent-gene summaries. **a**, Liver–LDL statistics for selected LR-context constituent-gene annotations. Four of 664 tested annotations are within-combination Bonferroni-positive; the lower band contains four illustrative lipid and extracellular contexts. Numbers beside Bonferroni-positive rows are one-sided empirical *P* values. **b**, Joint conditional S-LDSC *z* statistics for cell-intrinsic and spatial LR-gene annotations across LDL, triglycerides, CAD, T2D and BMI. Orange diamonds mark nominal-positive spatial LR-gene coefficients at one-sided *P* < 0.05, unadjusted across the 85 tissue–trait tests; relative cell-intrinsic and spatial LR-gene statistics do not classify a trait’s biological mechanism. **c**, Expression-derived sender and receiver summaries for five illustrative contexts in the cell-segmented liver data. Labels report the cell type with the largest ligand or receptor allocation, and bubble size increases with that allocation fraction. Sender–receiver orientation follows the curated LR label and is not genetically inferred. **d**, Constituent-gene annotation statistics for 668 liver–LDL contexts in UK Biobank and GLGC 2013 (*ρ* = 0.53). Diamonds identify three illustrative lipid contexts; the dashed line is the identity line. **e**, Aggregate spatial LR-gene statistics in standard Visium JBO001 and two cell-segmented Visium HD samples. The observed *z* statistics are 2.04, 1.93 and 1.63, respectively; the dashed line marks the nominal one-sided threshold *z* = 1.645. **f–h**, Expression-derived spatial LR-score maps for PCSK9–SORT1, LRPAP1–LRP1 and ANGPTL4–SDC4, displayed on a shared log_1_*p* colour scale. **i**, RGB overlay of the three normalized score maps. Context-level statistics in panels **a** and **d** use fixed constituent-gene annotations (§2.1).

Triglycerides were also above threshold (*z* = 2.08, *P* = 0.019), as was CAD (*z* = 1.96, *P* = 0.025); their high-ranked rows again included clearance-receptor genes. The two blood pressure traits in liver do not reach the aggregate threshold (SBP *z* = 0.92, *P* = 0.18; DBP *z* = 1.62, *P* = 0.052), so we do not use them as aggregate-supported liver case-study anchors; their empirically calibrated context-level results, including the liver–SBP rows, are reported in Supplementary Table 3, sheet “Liver”.

Expression-derived maps for top-ranked LR labels showed a 1.84-fold enrichment for spatial gradients along the portal–central axis (permutation *P* < 0.001; Extended Data Fig. 2f), consistent with the known zonation of hepatic lipid metabolism[35].

Consistent with the aggregate landscape (§2.3), T2D and BMI showed no edge signal in liver (Fig. 5b).

### 2.8 Trait-specific LR-context gene priorities in brain

Heart and liver illustrate tissue-specific pathway convergence; brain provides a within-tissue comparison of trait-specific edge-annotation profiles. In DLPFC, three traits showed positive aggregate edge associations using the same tissue data and LR database, yet each yielded a distinct set of top constituent-gene contexts (Fig. 6).

**Fig. 6.**
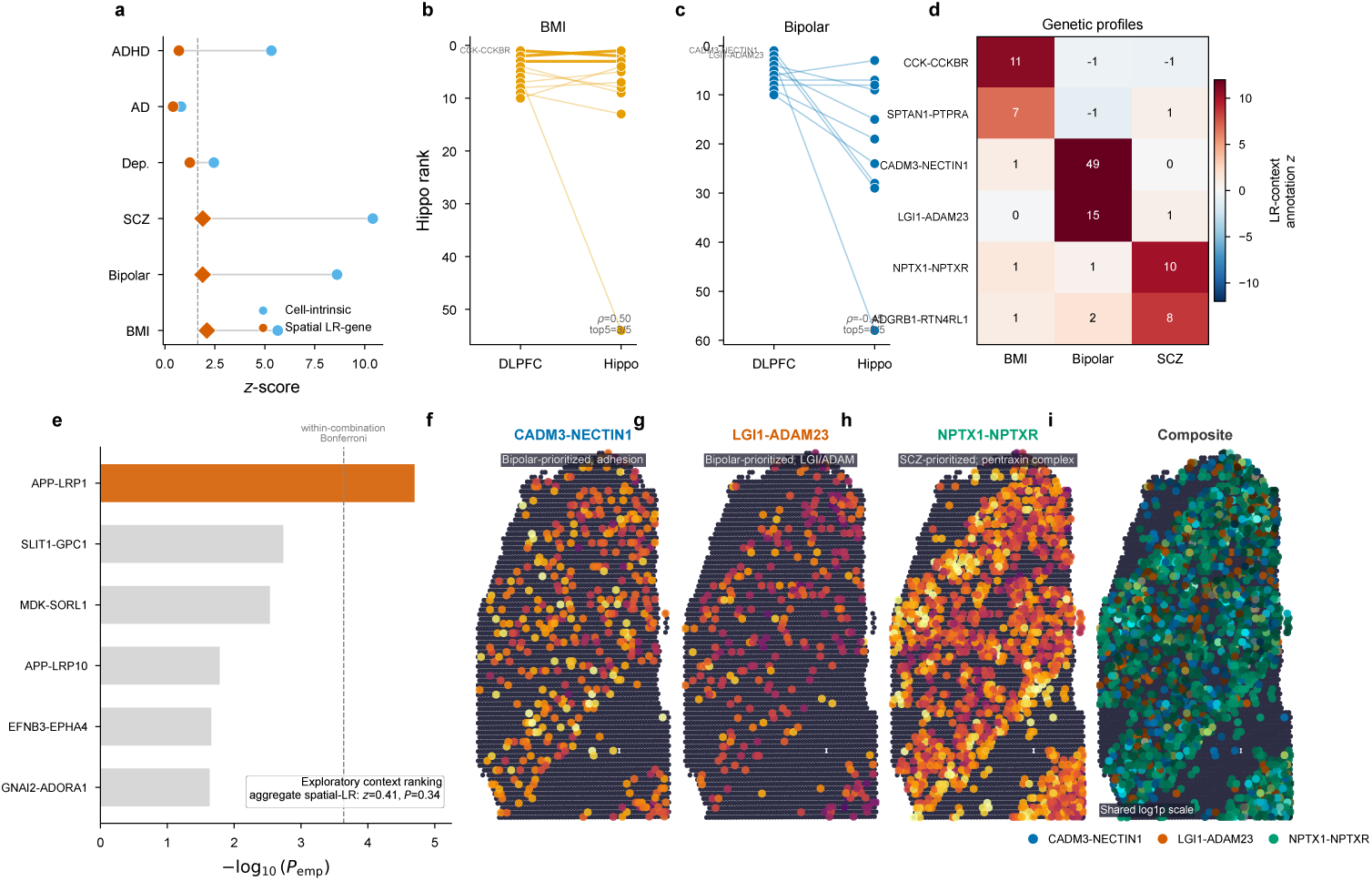
Trait-dependent constituent-gene annotation profiles in brain. **a**, Joint conditional S-LDSC *z* statistics for cell-intrinsic and spatial LR-gene annotations across six DLPFC trait analyses. Orange diamonds denote nominal-positive spatial LR-gene coefficients at one-sided *P* < 0.05, unadjusted across the 85 tissue–trait tests. **b,c**, Hippocampal ranks of the ten highest-ranked DLPFC constituent-gene annotations for BMI (**b**; *ρ* = 0.50, top-five overlap = 3*/*5) and bipolar disorder (**c**; *ρ* = *−*0.45, top-five overlap = 0*/*5). The rank correlations are descriptive and are calculated within the DLPFC-selected ten-row sets. **d**, Cross-trait *z* profiles of the two highest-ranked annotations selected separately for BMI, bipolar disorder and schizophrenia. Because rows were selected using the displayed traits, the diagonal concentration is descriptive rather than a formal test of trait specificity. **e**, The aggregate spatial LR-gene coefficient for DLPFC–AD is null (*z* = 0.41, *P* = 0.34). Among 218 context annotations, APP–LRP1 has the smallest one-sided empirical *P* value and reaches the within-combination Bonferroni threshold; the panel is therefore a constituent-gene ranking within a null aggregate. **f–h**, Expression-derived spatial LR-score maps for CADM3–NECTIN1, LGI1–ADAM23 and NPTX1–NPTXR, displayed on a shared log_1_*p* scale. **i**, RGB overlay of the three normalized maps. Context-level statistics use fixed constituent-gene annotations (§2.1).

#### BMI

BMI showed a positive aggregate edge association (*z* = 2.10, *P* = 0.018); the leading ranks included CCK–CCKBR, a CCKergic neuropeptide receptor axis linked to feeding/satiety and broader CNS neuromodulatory functions[36], and SEMA6B– PLXNA4, implicating axon guidance.

#### Bipolar disorder: trans-synaptic adhesion

Bipolar disorder and schizophrenia showed comparable positive aggregate edge signals (*z* = 1.89, *P* = 0.030 and *z* = 1.89, *P* = 0.029), yet their context rankings share no row. For bipolar disorder the CADM3– NECTIN1 context reaches FDR < 0.10 (*P*_emp_ = 3.4 × 10^−4^, FDR = 0.074; Supplementary Table 3, sheet “Brain DLPFC”). CADM3 (SynCAM3/Necl-1) is a neural adhesion molecule that localizes to non-junctional contact sites of presynaptic terminals and glial processes and trans-interacts with nectin-1[37]. The next-ranked context, LGI1–ADAM23, lies in the LGI/ADAM epilepsy-associated adhesion family: secreted LGI1 binds ADAM23, and altered ADAM23-dependent neuronal morphology contributes to lateral temporal epilepsy[38]; related LGI1–ADAM22 complexes organize trans-synaptic nanoalignment for synaptic transmission and epilepsy prevention[39]. These rankings prioritize synaptic and axon–glia membrane genes.

#### Schizophrenia

Schizophrenia’s top contexts differ from those for bipolar disorder and centre on excitatory synapse regulation: NPTX1–NPTXR, part of the neuronal pentraxin complex that recruits GluA4-containing AMPA receptors, with related NPTX2–NPTXR perturbation shaping excitatory drive onto fast-spiking parvalbumin interneurons[40, 41]; ADGRB1–RTN4RL1, pairing a BAI-family adhesion G-protein-coupled receptor with a Nogo-receptor-like partner; and the SIRPA–CD47 row, which corresponds biologically to CD47–SIRP*α* signalling between synapses/neurons and microglia, a brake on microglial pruning[42]. No schizophrenia context reached FDR < 0.10 after the correction described in §4.8, so these are rankings rather than formal findings.

The three traits prioritize different constituent-gene families in the same tissue against the same LR database—CCKergic neuropeptide biology for BMI, contact-dependent adhesion for bipolar disorder, and excitatory synapse regulation for schizophrenia. This trait-associated variation is inconsistent with a single invariant tissue ranking. The pentraxin and pruning-related genes for schizophrenia are consistent with the excitation–inhibition imbalance and synaptic-pruning accounts of the disorder, and the bipolar priorities with the myelin and oligodendrocyte changes reported in bipolar cortex[43].

The trait specificity of brain edge signals also extended across brain regions: in hippocampus, BMI was also nominal-positive (*z* = 2.37, *P* = 8.9 × 10^−3^) with CCK–CCKBR again among the top ranked contexts (Supplementary Table 3, sheet “Hippocampus”; Extended Data Fig. 1). The empirical null was run for heart, DLPFC, liver and gut, so hippocampal context-level values support ranking only; no calibrated individual-context claim is made for this tissue. Depression (*z* = 1.24, *P* = 0.11) and Alzheimer’s disease (*z* = 0.41) did not reach aggregate significance. For AD, context-level analysis prioritized the APP–LRP1 constituent-gene annotation at the within-combination Bonferroni threshold (*P*_emp_ = 2.0 × 10^−5^), motivating an APP/A*β*–LRP1 trafficking-clearance hypothesis despite the null aggregate. APOE– LRP1 had empirical *P* = 0.43 in the same ranking (Supplementary Table 3, sheet “Brain DLPFC”). The weaker common-variant signal (mean *χ*^2^ = 1.26 vs. 2.12 for SCZ) further limits aggregate sensitivity.

### 2.9 Biological content of prioritized constituent genes

If prioritized constituent genes provide complementary gene-level context, they should partially overlap with—but not recapitulate—existing gene-level annotations. We tested this prediction using two external reference annotations (drug target status, Mendelian disease genes) and a competitive gene-set test (Extended Data Fig. 5c–e). The aggregate analysis yielded a 13-combination nominal-positive pattern, with no individual panel-wide FDR < 0.05 result. Within-combination priority rows yield 31 unique constituent genes across all tiers (29 from tier 1 and tier 2 extracellular-verified contexts; Methods).

#### Drug target overlap

Of 29 tier 1+2 prioritized constituent genes, 17 (58.6%) have approved-drug interaction claims in DGIdb[44] (OR = 1.82, *P* = 0.08 versus the LR background of 44.0%; Extended Data Fig. 5c). The enrichment does not reach significance (*N* = 29), so the overlap provides contextual anchoring rather than validation. FLT1 (heart CAD, nominated receptor-side by the VEGFD–FLT1 row) marks a VEGFR1/sFLT1 component of VEGF-pathway-associated hypertension biology[45]; ERBB2 is the drug-targeted receptor named by the heart AD HLA-A–ERBB2 row and HER2 blockade can cause cardiotoxicity by disrupting cardiac NRG-1/ErbB2/ErbB4 survival signalling[46]; and LRP1 appears repeatedly as an endocytic/clearance-receptor axis across brain, heart and liver. Because several LRP1-containing rows share receptor-side structural support, we treat this recurrence as receptor-axis prioritization rather than as eight independent ligand-specific discoveries (Supplementary Table 9).

#### Mendelian convergence

One of 29 tier 1+2 prioritized constituent genes matched an OMIM Mendelian gene for the corresponding trait: APP for Alzheimer’s disease (OR = 0.75, *P* = 0.74). The rate is no higher than the ligand–receptor background, consistent with partial, trait-specific convergence rather than class-wide enrichment between monogenic causal genes and polygenic spatial LR-gene annotations. APP, a causal gene for rare familial early-onset Alzheimer’s disease, appears in the APP–LRP1 brain context at the within-combination Bonferroni threshold (*P*_emp_ = 2.0 × 10^−5^), with both constituent genes contributing SNP support.

#### Competitive gene-set test

Aggregating GWAS *P* -values to gene-level statistics (minimum *P* within ±10 kb, analogous to MAGMA[21]), prioritized constituent genes carried stronger associations than the LR background in 5 of 8 testable trait– tissue combinations containing priority rows (Wilcoxon rank-sum *P* < 0.05), with the strongest signals in liver–LDL (*P* = 1.1 × 10^−4^), liver–SBP (*P* = 1.3 × 10^−3^) and liver–TG (*P* = 3.4 × 10^−3^). This supports above-background GWAS concentration within the LR database, while remaining an internal consistency check because it shares GWAS input with EdgeMap (Methods).

Despite the statistical cost of testing sparse LR-context annotations (§4.8), fixed constituent-gene annotation profiles show concordance across independent GWAS cohorts and calibration methods (Extended Data Fig. 4b–e). Agreement across sections of the same tissue is not included here: as shown in §2.5, positive-scalar invariance reproduces it. Clustering context-level *z*-score profiles across traits organized these constituent-gene contexts into recurrent modules (Extended Data Fig. 2): a BMI-loaded neuropeptide/adhesion module anchored by CCK–CCKBR and SPTAN1–PTPRA, a bipolar-loaded adhesion/calcium-signalling module anchored by CADM3–NECTIN1 and LGI1–ADAM23, and blood-pressure modules including HSPG2–FGFR1 and CALR–SCARF1—a descriptive organization of recurring gene-set profiles rather than evidence for resolved molecular channels. These comparisons assess gene-set stability, not LR identity or direction.

#### Complementary fine-mapping analysis

As a complementary gene-level analysis, we tested whether genes with high edge specificity carry elevated fine-mapping signal from SuSiE/SuSiE-inf credible sets (Open Targets Genetics; Methods). Among nominal-positive aggregate case-study combinations with sufficient fine-mapping data, edge-annotated genes show 1.2–3.1-fold elevated PIP burden with nominal, unadjusted *P* < 0.05 in 7 of 10 testable trait–tissue combinations (Mann–Whitney), whereas the corresponding node comparison has *P >* 0.10 in all 15 testable combinations (Methods). At established GWAS loci, five of seven established locus genes—PCSK9, LDLR, APOB (LDL), LPL (TG), and AGT (SBP)—rank first by CSS among all genes within ±500 kb, showing that edge annotation can prioritize established locus genes (Extended Data Fig. 7h). Gene-level fine-mapping enrichment is consistent across Open Targets studies: for BMI, the edge-specific Spearman correlation between two studies reaches *ρ* = 0.98 (Extended Data Fig. 7i), supporting cross-study consistency of the fine-mapping signal. Because Open Targets studies can share participants, this comparison is not treated as independent-cohort replication. To address measured gene-level covariates (gene length, cis-SNP count, LD score and expression), we performed a matched competitive test restricting the universe to LR genes and matching each prioritized LR gene to up to four control-LR genes by Mahalanobis distance on these covariates. Within this matched background the association is attenuated but not eliminated: among 14 attempted comparisons, 11 have non-zero edge or control PIP burden and four have nominal *P* < 0.05 (liver–DBP, hippocampus–BMI, liver– CAD and liver–SBP; Extended Data Fig. 7e; Methods). This matched analysis is descriptive rather than definitive: part of the fine-mapping pattern persists within an LR-only, covariate-matched background, while the attenuation indicates that baseline gene properties contribute substantially.

## 3 Discussion

Across 17 traits and five human tissues, spatially weighted LR-gene annotations carry conditional heritability association that concentrates in biologically matched tissues: all 15 traits with both tissue classes rank the matched tissue higher (sign *P* = 3.1 × 10^−5^), and 11 of 13 nominal-positive combinations fall inside the prespecified matched set (*P* = 1 × 10^−4^). This pattern is stable across donors, receives support from independent GWAS analyses, and organizes into tissue-specific pathway families.

Prespecified and additional sensitivity analyses address broad LR-gene membership, spatial cell-type composition and several modeled null architectures; their convergence narrows these specific alternative explanations but does not exclude unmeasured annotation structure or residual confounding. A secondary analysis uses active LR contexts to prioritize their constituent genes, revealing largely non-overlapping candidates compared with standard gene-level methods.

Existing heritability methods—S-LDSC[2], scDRS[5], gsMap[6]—identify disease-relevant tissues and cell types but score each cell independently; EdgeMap adds a complementary spatially informed annotation layer. In heart, for example, the top node genes are sarcomeric and structural (TTN, MYH7, ACTA1), whereas the prioritized constituent genes GPC3, LRP1, VEGFD and FLT1 encode intercellular signaling molecules and all fall below the median of the same GSS distribution (ranks 8,900, 9,804, 12,608 and 10,087 of 15,217; §2.4).

A recurring observation across tissues is that edge-annotation and constituent-gene priorities organize into pathway families: vascular and extracellular matrix receptor axes in heart, lipoprotein receptors in liver, and trans-synaptic adhesion and synapse-regulation genes in brain. The structural audit helps calibrate this interpretation. In brain, APP–LRP1 and CADM3–NECTIN1 have mappable constituent genes on both sides and motivate follow-up. In heart and liver, several rows share receptor-side support, so their strongest use is to prioritize clearance-receptor axes and tissue localization. This reading suggests recurrent spatial LR-gene modules (Extended Data Fig. 2), with biologically related traits sharing correlated edge profiles within tissues (Extended Data Fig. 2c,e).

The controls described in §2.3–2.4 (cell-type composition, gene-level cis-LD burden and expression, LR specificity, locus ablation; Fig. 3c,d,f) indicate that the edge signal is not reducible to a single confounding mechanism. The LR-database shuffle is a database-specificity diagnostic rather than a requirement for every tissue to pass: it asks whether the curated LR resource outperforms expression-matched LR-like databases. Its significance in liver and gut supports database-specific LR organization there; the nonsignificant heart and brain results suggest that those aggregate edge signals may reflect broader spatial organization of cell-surface and receptor-axis genes shared by expression-matched alternatives. Node and edge *z*-scores are positively correlated across the landscape (Spearman *ρ* = 0.66), as expected because both annotations are strongest in trait-relevant tissues, but it does not make the two layers redundant: the edge annotation prioritizes largely non-overlapping genes (Extended Data Fig. 3a), remains coherent in within-trait matched-tissue contrasts, and shows complementary fine-mapping concordance. The edge layer is complementary to, not independent of, cell-intrinsic tissue relevance.

A controlled Cross-Moran specification that conditions on LR membership and on ligand- and receptor-side marginal expression provides an additional sensitivity analysis, with two nominal-positive contrasts and greater coordinate-shuffle attenuation for the controlled term than for the same edge statistic without membership terms within the targeted, non-exhaustive 28-combination sensitivity panel (median 30.5% versus 2.9% among the 10 contexts with controlled *z >* 1; §4.32). We treat this as supportive sensitivity evidence rather than as a primary validation layer because attenuation is heterogeneous and proportional changes are unstable near zero.

Taken together, the evidence is not driven by a single test: within-trait tissue coherence, landscape-wide concentration under a correlated null, donor-level aggregate replication, independent-GWAS support and cell-segmented Visium HD collectively support an association between spatial LR-gene annotations and heritability.

Two observations carry broader implications. First, the brain analysis illustrates how the same tissue can carry trait-specific annotation signals rather than a generic neural enrichment. Bipolar disorder prioritizes trans-synaptic adhesion contexts including CADM3–NECTIN1 and LGI1–ADAM23, whereas the schizophrenia ranking emphasizes excitatory synapse organization through NPTX1–NPTXR (§2.8). Node-centric methods can identify individual genes in these pathways; EdgeMap asks whether spatially weighted LR-gene annotations add conditional association and then organizes gene priorities by active LR labels, a distinction not resolved by single-cell scoring. Second, expression-derived maps for prioritized LR labels are enriched for gradients along tissue axes (§2.7, §2.8; Extended Data Fig. 2f).

These properties—complementarity, trait specificity and spatial organization— suggest that EdgeMap can refine GWAS interpretation from “which tissues are relevant” to “which spatial LR-gene contexts merit mechanistic follow-up.” Current GWAS-to-drug pipelines prioritize cell-intrinsic targets, whereas intercellular interfaces include receptor, ion-channel, transporter and secreted-protein biology represented in approved target maps[47]. Drug-target catalogs are also historically shaped by tractability and therapeutic attention, so any overlap with prioritized constituent genes should be read as contextual prioritization rather than validation. That 16 of 31 unique prioritized constituent genes (52%) are absent from standard gene-level prioritization (§2.4; Extended Data Fig. 5) suggests that LR-context organization can complement conventional gene-level candidate lists; it does not by itself confer causal or relation-level support.

Several limitations should be noted. First, individual trait–tissue *z*-scores are moderate: the 13 nominal-positive combinations span *z* = 1.77–2.37, as expected when aggregate annotations carry limited per-SNP variance in the S-LDSC framework[2]. Computed from the standard errors the analysis produced rather than from the power simulation, which overstates precision by 4.9-fold in the median-SE comparison, 80% average power requires *τ*_edge_ ≈ 6.7 × 10^−9^, so most real trait–tissue combinations sit below the detection boundary (Supplementary Table 2). The 13 of 85 threshold count is therefore a descriptive view of the landscape, not a discovery count or an estimate of the prevalence of spatial LR-gene association. Interpretation instead rests on the full-panel coherence tests and their stated uncertainty.

Second, LR-context annotations are sparse, so block-jackknife standard errors lose *N* (0, 1) validity (§4.8). Formal 50,000-replicate empirical calibration was completed for 18 of 85 tissue–trait settings. Moreover, for context *p* the tested annotation is **a***_p_* = *s_p_***m***_p_*: positive scalar scaling leaves *z*-scores and *P* -values unchanged, and the context’s own membership covariate is exactly collinear. The context-level analysis therefore prioritizes constituent genes within CSS-active LR contexts but cannot identify LR relations, direction or interaction effects. This is a design boundary, not a limitation that additional data can overcome within the current estimand. Third, as noted in §2.1, the LIANA Consensus resource includes pairs whose partners lack documented extracellular localization (e.g., calmodulin–channel and spectrin– phosphatase interactions). All context-level biological interpretation in this paper uses extracellular-verified pairs (tiers 1 and 2, ~90% of pairs; §4.35). Under this default filter, 12 of 13 originally nominal-positive combinations remained nominal-positive; the one that became marginal was brain–bipolar disorder (*P* = 0.109), consistent with the mixed synaptic and intracellular nature of adhesion signaling. A stricter secreted-ligand membrane-receptor filter (tier 1 only; 54% pair retention) retained 7 of 13, with robust retention in heart and liver. In brain, the context-level rankings after filtering converge on trans-synaptic adhesion (CADM3–NECTIN1, LGI1–ADAM23) and excitatory synapse regulation (NPTX1–NPTXR), which constitute the biological interpretation presented in §2.8.

Fourth, Visium’s ~55 *µ*m resolution means each spot contains 1–10 cells; the “edges” therefore reflect communication between local cell populations rather than individual cell–cell contacts. Cross-resolution support from cell-segmented Visium HD data (median nearest-neighbour distance ~13 *µ*m; §2.5) supported the liver LDL edge signal, and sender–receiver analysis at 8 *µ*m resolution showed that all five highlighted liver–LDL maps allocate ligand- and receptor-side expression to distinct cell types (e.g., hepatocyte F9 and stellate-cell LRP1). This supports spatial localization consistent with intercellular geometry. In heart, Kuppe deconvolution provides compartment-level localization for selected cardiovascular contexts; single-cell platforms (MERFISH, Slide-seq) may further sharpen spatial resolution. Fifth, the aggregate edge test for Alzheimer’s disease was null in DLPFC. APP–LRP1 is therefore a context-level priority that motivates trafficking-clearance follow-up but does not by itself validate the relation. Relation-level resolution would require designs that distinguish joint from constituent-gene effects and incorporate direction- or perturbation-specific information. Finally, although the complementary fine-mapping analysis—including matched competitive testing against measured-covariate-matched LR controls—shows nominally elevated fine-mapping signal in some edge-annotated gene sets (§2.9), EdgeMap does not itself identify causal variants; integration with spatial causal mediation remains a natural next step.

Looking ahead, the joint node–edge analysis suggests a useful division of labour: the statistical model tests aggregate conditional association and prioritizes constituent genes, while the spatial LR labels formulate mechanistic hypotheses for experiments. For example, CADM3–NECTIN1 and LGI1–ADAM23 organize bipolar-disorder priorities around adhesive contact biology. Conditional knockouts, adhesion-blocking experiments and future relational annotations that contrast joint with single-gene effects can test the missing relation-level estimand directly. As single-cell spatial technologies mature, extending the conditional-association framework established here to developmental stages, perturbations and relational annotations can turn bounded LR-context priorities into falsifiable experiments.

## 4 Methods

### 4.1 Spatial transcriptomics preprocessing

Spatial transcriptomics data were loaded from h5ad format and preprocessed using Scanpy[48]: genes expressed in fewer than 10 cells were removed, counts were librarysize normalized to 10,000 per cell, and log1p-transformed. Spatial coordinates were extracted from the obsm[’spatial’] field.

### 4.2 Spatial graph construction

We constructed a *k*-nearest-neighbor graph (*k* = 6) from spatial coordinates using Euclidean distance, where **p***_i_* R^2^ denotes the spatial coordinates of cell *i*. Edge weights follow a Gaussian kernel:

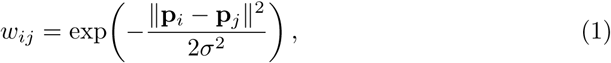

where *σ* = *d*_max_*/*3 and *d*_max_ is the maximum Euclidean distance for retaining a neighbor in the KNN graph (default 3,000 *µ*m for 10x Visium). Self-edges are excluded (*w_ii_* = 0) so that the LR activity proxy combines ligand expression in neighbouring spatial units with local receptor expression rather than using a same-unit product. The weight matrix is symmetrized: *W ←* (*W* + *W^⊤^*)*/*2.

### 4.3 Ligand–receptor database

We used the LIANA Consensus resource[7], which integrates curated LR pairs from CellChatDB[16], CellPhoneDB, ICELLNET[49], connectomeDB2020, and CellTalkDB[50], filtered by literature support and protein localization (4,624 interactions). An LR pair was retained if every subunit gene was expressed in at least 5% of spots.

### 4.4 Spatially weighted LR activity proxy

Using a mass-action-inspired expression product commonly adopted in cell–cell communication scoring[16], for each curated LR label *p* with ligand subunits *l_p_*and receptor subunits *r_p_*, we computed a per-cell LR activity proxy:

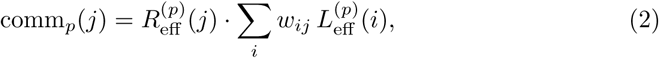

where 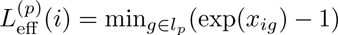 and 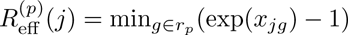 are effective expression values after reversing the log1p transform (*x_ig_* is the log1p-normalized expression of gene *g* in cell *i*). The min over subunits implements the standard bottleneck model for heteromeric complexes[17]. This score summarizes spatially weighted ligand–receptor expression; it does not measure molecular binding, signaling flux or diffusion.

### 4.5 Gene specificity score (GSS)

For each gene *g*, expression values were ranked across all *n* cells (one ranking per gene; ties broken arbitrarily). The gene-level GSS is the ratio of local to global geometric mean rank, maximized over all cells (max rather than mean, to capture the location where gene *g* is most spatially concentrated):

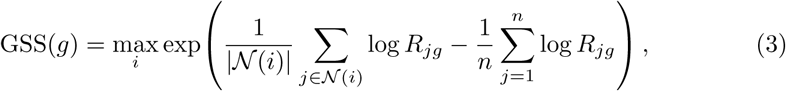

where *n* is the total number of cells, *R_jg_*is the rank of cell *j* among all *n* cells for gene *g* (ascending; rank 1 = lowest expression, including zeros), and (*i*) is the spatial neighborhood of cell *i* including *i* itself ((*i*) = *k* + 1 for interior cells).

The maximum is used here whereas CSS uses the 95th percentile, because the two quantities have different support: every gene is expressed in nearly every spot, so the maximum identifies the one neighborhood where a gene is most concentrated, while each LR pair is active in only a minority of spots, where a percentile is the more stable summary. To assess sensitivity to this choice we recomputed all 85 tissue–trait combinations with GSS aggregated at the 95th percentile instead: Spearman *ρ* = 0.941 for node and 0.988 for edge *z*-scores. No node result changes nominal-threshold status; three edge results change status, all in the direction of adding nominal-positive combinations under the 95th-percentile aggregation. Two are close to threshold, whereas liver–SBP reflects the aggregation choice (Supplementary Table 5, sheet “Max vs p95”).

### 4.6 Context specificity score (CSS)

For each LR label *p*, the context-level score is the 95th percentile of per-cell LR activity specificity: PairScore(*p*) = *Q*_95_(comm*_p_*(*j*)*/*com̄m*_p_*). The 95th percentile captures the spatial concentration of the expression-derived activity proxy while remaining robust to outliers (robust across the range 85–99%). Pairs with PairScore(*p*) ≤ 1 are set to zero. The gene-level CSS is the maximum pair score across all LR pairs in which gene *g* participates (max ensures that a gene’s CSS reflects its highest expression-derived LR-context score):

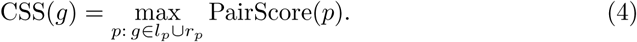

Max aggregation is an EdgeMap design choice: it ensures that a hub gene appearing in many pairs (e.g. ITGB1 in 456 pairs) receives the score of its highest expression-derived LR-context score rather than an inflated sum. As a sensitivity analysis we also computed CSS using mean-over-pairs aggregation; landscape-level results were highly concordant (Spearman *ρ* = 0.996, concordance of significance 83*/*85), indicating that the gene-level aggregation rule is not a critical degree of freedom. The exploratory context-level branch (§4.8) does not use the max-over-context aggregation. Its test statistic is invariant to the magnitude of any positive context score, although the same expression-derived score determines which contexts pass the activity gate.

### 4.7 EdgeMap joint regression

Gene-level scores were mapped to SNP-level annotation LD scores using the precomputed SNP–gene weight matrix **W** *∈* R*^M×G^* from gsMap[6], whose entry 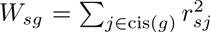 aggregates squared LD correlations over all SNPs *j* in the cis-window of gene *g*:

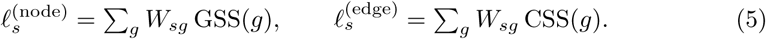

The joint regression extends the stratified LD score regression (S-LDSC) framework[2] by introducing these tissue-specific node and edge annotations alongside baseline annotations; the regression machinery and block-jackknife inference are unchanged. The model is:

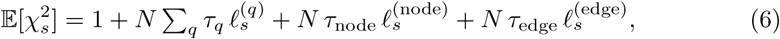

where 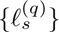 are baseline LD score annotations and *τ* coefficients are estimated by weighted least squares with block-jackknife standard errors over ~200 non-overlapping LD blocks. Blocks are formed from runs of consecutive SNPs, so the regression rows are sorted by chromosome and position before the jackknife: the row order that a merge onto summary statistics produces is not genomic for every GWAS, and blocks formed from it are genome-wide samples rather than regions. The point estimate is unaffected by the ordering, whereas the standard error can be affected (Supplementary Table 5, sheet “Jackknife Ordering”; Supplementary Table 1 summarizes the associated provenance and quality-control checks). We report one-sided *P* values for *τ*_edge_ *>* 0, as our primary hypothesis is directional: we test for heritability enrichment in the edge annotation rather than depletion.

### 4.8 Exploratory LR-context constituent-gene testing

For each LR context *p* with context specificity *s_p_ >* 1.0, let **m***_p_* denote the binary vector marking its constituent ligand and receptor genes. The gene annotation is **a***_p_* = *s_p_***m***_p_*, and its SNP annotation and LD-score vector are therefore the same scalar multiple of those obtained from **m***_p_*. Multiplying a regression column by the positive scalar *s_p_* rescales its coefficient and standard error by 1*/s_p_* but leaves its *z*-score and empirical or parametric *P* -value unchanged. The context’s own membership annotation is thus exactly collinear with the tested annotation. Context specificity gates which LR contexts enter this analysis, but does not weight their genetic test statistics. The three-annotation model (baseline + node + one LR context) therefore tests whether the context’s constituent-gene annotation has a positive conditional S-LDSC coefficient. It does not identify the ligand–receptor relation, its direction, a molecular interaction, or an interaction effect in the statistical sense.

Because context-level constituent-gene annotations are sparse (∼1,000 nonzero SNPs out of ~1,000,000), the block-jackknife standard errors that underlie S-LDSC *z*-scores are poorly calibrated: most of the ~200 LD blocks contain zero annotated SNPs and contribute no information, causing the null *z*-distribution to depart sharply from *N* (0, 1) (empirical sd ≈ 4.4, excess kurtosis 50–5,000, KS *P* < 10^−100^; Fig. 2b), with a spike-and-slab structure reflecting the sparse annotation geometry. This is a known limitation of S-LDSC for small annotations[3, 51]: the block-jackknife test does not control type-I error once an annotation overlaps too few genomic blocks, both because the jackknife estimate of the standard error is noisy at that scale and because the coefficient estimate is non-normal[51].

We therefore calibrate context-level tail probabilities by simulating 50,000 null GWAS for each tissue and retaining the resulting null distribution separately for every testable annotation. Formal empirical calibration was completed for 18 of the 85 tissue–trait settings. The calibrated subset comprised all 12 aggregate nominal-positive settings in brain, gut, heart and liver, plus six case-study or within-tissue comparators (brain AD and depression, gut CAD, and liver DBP, SBP and UC). Hippocampal nulls were not generated. This targeted, non-exhaustive subset was not sampled to estimate a discovery rate across settings; the remaining settings support aggregate analyses or rankings only; no study-wide discovery count is claimed. Under *H*_0_: *τ*_node_ = 0, *τ_p_* = 0 (a global null in which only baseline LD structure generates association signal), we simulated GWAS *χ*^2^ statistics from the real baseline LD scores, ran the full context-level S-LDSC pipeline for every testable annotation and recorded all null *z*-scores. Each annotation accumulates its own 50,000-element null distribution, shaped by that annotation’s sparsity and LD structure; the null is therefore annotation-specific, not pooled across contexts. The empirical *P* -value for each context annotation is

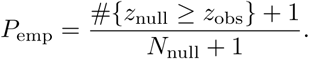

Because each annotation’s null *z*-distribution is shaped by its own annotation sparsity and is generally not centered at zero, the comparison is made against that annotation’s null rather than against *N* (0, 1). The empirical *P* -value measures the right tail; only rows with positive fitted coefficients are eligible for prioritization, although complete QC tables retain all tested rows. Context annotations whose observed *z* falls in the left tail—depletions relative to the annotation-specific expectation—are not prioritized. The hypothesis under test is that a constituent-gene annotation carries more per-SNP heritability than the baseline model predicts, not that it differs from the baseline in either direction; the stated alternative is therefore one-sided. Because positive node enrichment could in principle alter the context-level null through heteroskedasticity, we performed conditional-null sensitivity analyses across all eleven trait–tissue settings, simulating under each setting’s observed *τ*^_node_ (spanning a 130-fold range) with *τ_p_* = 0 (5,000 replicates each). Context-annotation null standard deviations were nearly identical (median SD ratio 0.97–1.02, overall mean 1.00), and empirical *P* - value ranks were near-perfectly concordant (Spearman *ρ* 0.998), supporting use of the global null for calibration (Extended Data Fig. 4d,e). Within-combination Bonferroni and Benjamini–Hochberg FDR summaries were applied to the empirical *P* -values across all testable context annotations within each tissue–trait combination. Annotations with null standard deviation < 0.5 were excluded as untestable (annotation too sparse). This empirical-null approach estimates tail probabilities from the simulated annotation-specific null rather than imposing a parametric *N* (0, 1) reference distribution, yielding a minimum empirical *P* of 2.0 × 10^−5^ from 50,000 null replicates with the add-one denominator 50,001. Although the LD-score vector for a testable context annotation typically contains ∼1,000 nonzero SNPs, those SNPs derive from cis windows around one or a few ligand/receptor genes and can be concentrated in a small number of genomic neighborhoods. Raw context-annotation *z*-scores can be very large (e.g., *z >* 30) when one such neighborhood overlaps a major GWAS locus; such values reflect instability of block-jackknife *z*-scores for sparse annotations rather than proportionally large heritability contributions. The empirical *P* -value compares the observed *z* against the same annotation’s null distribution under the specified simulation model and is the inferential quantity used for this exploratory calibration.

### 4.9 LR-context structural support audit

Empirical calibration assigns a tail probability for a fixed annotation, but it does not by itself determine which constituent genes contribute SNP support. We therefore audited every reported context row for structural support. For each tissue–trait regression frame, after GWAS, baseline and regression-weight SNPs were intersected and sorted in genomic order, we counted: (i) which ligand and receptor genes were present in the autosomal SNP–gene weight matrix and contributed at least one nonzero SNP; (ii) the number of supported SNPs on the ligand side, receptor side and their union; and (iii) the number of delete-one-block jackknife blocks represented by each side and by the union. We also flagged pairs whose contributing autosomal gene set was identical to another pair in the same tissue–trait combination.

This audit is a structural-support and sparsity check, not evidence for an LR relation and not an additional significance filter. A context was flagged as single-sided if either the ligand or receptor side had zero supported blocks, as single-block if the ligand–receptor union occupied at most one jackknife block, and as non-unique if another pair in the same combination mapped to the same contributing gene set. Flagged rows can still contribute to gene-level or receptor/locus prioritization, but they should not be overinterpreted as identifying a two-gene communication relation. Even rows with both constituent genes mappable remain gene-set contexts because the tested annotation contains no relational or interaction term. Complete results are reported in Supplementary Table 9.

#### Null *χ*^2^ generation

Under *H*_0_ each null *χ*^2^ statistic is generated as

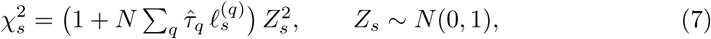

where *{τ*^*_q_}* are baseline coefficients estimated by S-LDSC from the observed GWAS summary statistics and 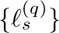 are the baseline annotation LD scores. Each LD score 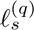 aggregates pairwise LD between SNP *s* and all other SNPs weighted by annotation *q*, so the mean structure of the synthetic *χ*^2^ reflects the genome’s LD architecture.

Although the noise terms *Z_s_* are drawn independently, the S-LDSC block jackknife is robust to within-block LD correlation, as confirmed by the LD-aware validation below. The trait-specific heritability enters through *τ*^*_q_*, which is estimated from the data rather than fixed at any pre-specified value; the fixed *h*^2^ = 0.3 mentioned in the simulation study (§2.2) applies exclusively to the power analysis (Fig. 2), not to real-data inference. As a robustness check, we tested heavy-tailed effect-size distributions (*t*_3_, *t*_5_, Laplace, point-normal mixture) within the genotype-derived null battery (see below) and confirmed that context-annotation null calibration is insensitive to heavy tails (type-I error 4.3–5.1% across all distributions; Extended Data Fig. 7d). To directly test whether LD-induced correlations between SNPs affect null calibration, we implemented an LD-aware null in which noise terms are drawn jointly from (**0**, **R***_b_*) within each of 1,702 LDetect blocks, where **R***_b_* is the empirical LD correlation matrix (Cholesky decomposition of plink LD estimates among HapMap3 SNPs). At the aggregate level, the LD-aware null yields edge type-I error of 4.4% (1,000 replicates; sd(*z*) = 1.04; KS uniform *P* = 0.08), identical to the independent null. At the context-annotation level, the SD ratio between LD-aware and independent nulls has median = 1.01 (IQR [0.87, 1.19]; 370 pairs, 2,000 replicates each; Extended Data Fig. 7g), indicating that residual within-block LD correlation has little practical effect on calibration in this validation setting. We do not extrapolate this diagnostic beyond the tested annotations and simulation specifications.

#### Genotype-derived null calibration

To provide the most stringent validation using individual-level genotype data, we conducted a genotype-derived null battery using 1000 Genomes EUR Phase 3 (503 unrelated individuals, QC’ed to HapMap3 SNPs). For each replicate, causal variants are drawn with probability *π* under each architecture and assigned variance-normalised effect sizes from a specified distribution (Gaussian, *t*_3_, *t*_5_, Laplace, or 90/10 point-normal mixture). Marginal associations are computed as 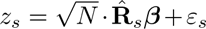, where **R**^^^ is the sample LD matrix from the 503 individuals, producing GWAS summary statistics with realistic LD structure. We tested 22 configurations: six core architectures (sparse *π* = 0.001, 0.01, 0.1; infinitesimal *π* = 1; 5× node-enriched with *τ*_edge_ = 0; 5× LR-membership-enriched with context-specific enrichment = 0) across three representative traits with published heritability and sample size (SBP: *h*^2^ = 0.15, *N* = 340k; LDL: *h*^2^ = 0.09, *N* = 344k; SCZ: *h*^2^ = 0.24, *N* = 77k), plus four heavy-tailed distributions at *π* = 0.01 (SBP only). Each configuration was run for 1,000 replicates through the full EdgeMap S-LDSC pipeline (aggregate regression and context-level FWL block-jackknife across all 370 active LR contexts). Context-annotation type-I error at *α* = 0.05 was 4.2–5.3% across all uniform and node-enriched architectures (Supplementary Table 6, sheet “Genotype Null Battery”; Extended Data Fig. 7d).

### 4.10 Pattern-level significance

We first applied Benjamini–Hochberg correction across all 85 aggregate tests; no combination reached panel-wide FDR < 0.05 (*q*_min_ = 0.21). Separately, three complementary pattern-level tests ask whether the nominal *P* < 0.05 crossings and signed *z*-scores are coherently concentrated in biologically matched tissues. These tests do not convert individual nominal results into adjusted discoveries. (1) A one-sided binomial test comparing the number of nominal-positive combinations (*k* = 13 of *n* = 85 at *α* = 0.05) against the null rate of 5%. (2) A trait-label permutation test for biological coherence. Biologically matched combinations were defined by the broad biological rule used in the Results and applied uniformly across the panel: heart with SBP, DBP, CAD, AF, heart rate ; brain DLPFC and hippocampus with SCZ, bipolar, depression, ADHD, BMI ; liver with LDL, TG, T2D, BMI, CAD, SBP, DBP ; gut with UC, Crohn’s . This covers 24 of the 85 combinations. A narrower 14-pairing enumeration is reported as a sensitivity analysis; because it omits ten combinations that the rule covers and all ten are null, we report the rule-based figures in the Results and both versions in Supplementary Table 5. Trait labels were shuffled within each tissue 10^4^ times, preserving each tissue’s edge-*z* distribution, and two statistics were compared to their null distributions: the count of nominal-positive results falling in biologically matched combinations, and the mean edge *z*-score difference between matched and unmatched combinations. (3) A Mann–Whitney *U* test comparing edge *z*-scores in biologically matched combinations versus all others.

The binomial test in (1) treats the 85 combinations as independent, whereas combinations sharing a tissue share an annotation and combinations sharing a trait share a GWAS. We therefore repeated it against a null that carries this dependence. Writin*_√_*g *z_tr_* for *_√_* tissue *t_√_*and trait *r*, we fitted the exchangeable two-way structure 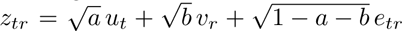 with independent standard normal *u*, *v* and *e*, so that *a* is the correlation between combinations sharing a tissue and *b* that between combinations sharing a trait. A full 17 × 17 trait correlation matrix is not estimable from five tissues; the two-parameter form is identified by the row and column mean squares of the observed *z*-score matrix. Estimates were *a* = 0.068 and *b* = 0.101. Drawing 2 10^5^ null matrices under this structure gave *P* = 2.7 × 10^−3^ against 13 observed, where the naive binomial gives 3.0 × 10^−4^. Accounting for shared tissue and trait structure therefore changes the estimate by about an order of magnitude while retaining evidence for a panel-level excess. Because the variance components are estimated from the observed matrix, this calibrated *P* value remains model-dependent and is reported with that limitation.

### 4.11 Standardised effect size and enrichment

Because *τ* coefficients are not comparable across annotations of different size, we also report the standardised effect size *τ^∗^* = *τ_c_ ×* sd(*a_c_*) × *M/h_g_*^2^ 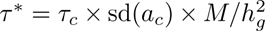 following Gazal et al.[52], where sd(*a_c_*) is the standard deviation of the raw annotation across SNPs, *M* the number of SNPs and 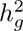 the SNP heritability from the joint model, together with the per-annotation enrichment (proportion of heritability divided by proportion of SNPs). Raw annotations were recomputed from the SNP–gene cis-indicator files. For each tissue we combined *τ^∗^* across the 17 traits by DerSimonian–Laird random-effects meta-analysis, reporting *I*^2^ as a heterogeneity summary. Values for all 85 combinations and the per-tissue meta-analyses are given in Supplementary Table 2.

We also examined the distribution of *P* -values directly: quantile plots for the aggregate tests and for the context-annotation empirical *P* -values are shown in Extended Data Fig. 6a,b. The context-level panel plots the empirical values on which exploratory prioritization rests, not parametric *z*-based values, and its bulk lies on the null diagonal (*λ*_GC_ = 1.03), which is the calibration the empirical-null procedure is designed to deliver.

### 4.12 Prioritized constituent-gene definition

For the downstream descriptive analyses below, *prioritized constituent genes* are genes in at least one LR-labelled annotation reaching within-combination FDR < 0.10 under empirical-null calibration (§4.8); genes appearing in multiple contexts are counted once per tissue–trait combination. This convenience set is not a study-wide discovery set and does not imply that the named LR relation is genetically identified. The LR database lists a few complexes in both orientations—for example NRXN1–NLGN2 and NLGN2–NRXN1—and the edge statistic is symmetric in its two partners, so the two orientations encode the same gene-membership annotation: across the 6,932 rows the 35 reverse orientations agree on *z* and empirical *P* to machine precision. We retain both source rows in Supplementary Table 3, flag the duplicate orientation and count its constituent genes once. Biological-context analyses (drug target overlap, Mendelian overlap) restrict prioritized genes to extracellular-verified contexts (tiers 1 and 2; §4.35), yielding 29 unique genes after pooling across nominal-positive aggregate combinations. Statistical analyses that assess GWAS signal strength (competitive gene-set test, gene-level method comparison) use all 31 prioritized constituent genes regardless of tier. All comparisons use the 1,877-gene LIANA Consensus LR database as the background set and are interpreted as gene-level summaries.

### 4.13 Drug target overlap analysis

Approved-drug-associated genes were obtained from DGIdb v4[44] via the GraphQL API, retaining genes with at least one interaction classified as having an approved drug (3,439 genes). Enrichment was assessed by one-sided Fisher’s exact test comparing the fraction of approved-drug-associated genes among prioritized constituent genes to the background rate (826/1,877 = 44.0%). A pooled analysis combined unique prioritized genes across all nominal-positive aggregate combinations; a sensitivity analysis using all prioritized genes regardless of tier is reported in Supplementary Table 4, and the gene-by-gene tier assignment in the “Prioritized Genes” sheet of Supplementary Table 4.

### 4.14 Mendelian gene enrichment

Known causal genes for Mendelian forms of 15 complex traits were curated from OMIM. For each nominal-positive aggregate trait–tissue combination, we tested whether tier 1+2 prioritized constituent genes are enriched for the corresponding Mendelian genes using a one-sided Fisher’s exact test. A pooled analysis combined unique prioritized genes across all nominal-positive aggregate combinations.

### 4.15 Locus-to-Gene enrichment

Locus-to-Gene (L2G) predictions were obtained from the Open Targets Platform (release 25.03)[22]. For each trait, we identified L2G-prioritized genes (score ≥ 0.5) from GWAS studies matching the trait by EFO disease identifiers and keyword search. Enrichment of prioritized constituent genes among L2G-prioritized genes was tested by one-sided Fisher’s exact test against the LR database background, both per trait–tissue combination and pooled.

### 4.16 Competitive gene-set enrichment

To test whether prioritized constituent genes carry stronger GWAS associations than expected from the LR database background, we performed a competitive gene-set test analogous to MAGMA[21]. For each trait, SNP-level *P* -values were mapped to genes using a *±*10 kb window, retaining the minimum *P* -value per gene. Gene-level *P* -values were log_10_-transformed to obtain association scores. For each trait–tissue combination with within-combination empirical FDR < 0.10 contexts and at least two mappable prioritized genes, we compared the association scores of prioritized genes (all tiers) against all other genes in the LIANA Consensus LR database using a one-sided Wilcoxon rank-sum test. Combinations with fewer than two mappable prioritized genes were not eligible for the rank-sum comparison.

### 4.17 Gene-level method comparison

To quantify the complementarity of EdgeMap with existing gene prioritization methods, we compared prioritized constituent genes (all tiers) against two standard approaches for each of the 13 nominal-positive aggregate trait–tissue combinations. (1) *MAGMA-like gene-level testing:* for each trait, SNP-level *P* -values were mapped to genes within a ±10 kb window using the same procedure as the competitive gene-set test above; genes reaching Bonferroni significance (*P* < 2.5 × 10^−6^, correcting for ~20,000 protein-coding genes) were classified as MAGMA-significant. Gene coordinates were obtained from Ensembl GRCh37 (release 87) and SNP positions from the 1000 Genomes EUR phase 3 reference panel. (2) *Open Targets L2G:* for each trait, L2G-prioritized genes (score ≥ 0.5) were obtained using per-trait EFO and keyword matching as described above, restricted to studies of the *same* trait (i.e., per-trait matching rather than pooled across all traits as in the enrichment analysis). A prioritized gene was classified as “found by MAGMA or L2G” if it appeared in either method’s gene list for the specific trait in which it was identified by the LR-context analysis. The unique fraction was computed as the proportion of prioritized genes not found by either method.

### 4.18 Complementary fine-mapped variant analysis

As a complementary analysis of the same GWAS evidence, we analysed SuSiE and SuSiE-inf credible sets from Open Targets Genetics (release 25.03). All variants with posterior inclusion probability (PIP) *>* 0.01 were extracted for each trait, matched by EFO disease identifiers and keyword search. Variants were mapped to genes via cis-regulatory windows derived from SNP–gene pair files (the same windows used to construct the SNP–gene weight matrix).

We performed five descriptive layers of analysis; nominal *P* -values were not adjusted across layers or tissue–trait comparisons. (1) *Gene-level PIP burden:* pergene PIP sums were computed and genes in the top 20% of edge specificity compared against the rest (one-sided Mann–Whitney *U*). (2) *Variant-level cis enrichment:* for each variant, we determined whether it maps to at least one gene with CSS in the top 20%; a logistic regression of *P* (in edge-cis) on log_10_(PIP) tested the association between fine-mapping confidence and edge-gene proximity. (3) *Locus-level contextualization:* for established GWAS loci (PCSK9, LDLR, APOB, LRP1, SORT1 for LDL; LPL for TG; AGT for SBP; and others), all genes within ±500 kb of the established locus gene were ranked by CSS. (4) *Cross-study consistency:* for traits with multiple GWAS studies in Open Targets, gene-level PIP sums were computed per study and the Spearman correlation was calculated across studies, both genome-wide and restricted to edge-annotated genes. (5) *Matched competitive test:* to address measured gene-level covariates, we restricted the gene universe to LR genes (genes appearing in ≥1 active ligand–receptor pair) and classified the top 20% by edge specificity as edge-LR, the remainder as control-LR. Each edge-LR gene was matched without replacement to up to four control-LR genes by Mahalanobis distance on four covariates: log(gene length), log(*n*_cis-SNPs_), mean expression, and mean LD score. LR degree (connectivity) was not matched because it contributes to CSS construction; residual degree-related differences therefore remain possible. Gene-level PIP burden and credible-set gene overlap were compared between edge-LR and matched control-LR genes (one-sided Mann– Whitney *U* and Fisher’s exact test, respectively). Matching balance was assessed by standardised mean difference (SMD) on each covariate.

### 4.19 Simulation study

We generated synthetic GWAS *χ*^2^ under controlled architectures using real annotation LD scores from spatial transcriptomics. Expected 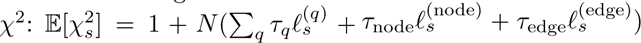, where baseline coefficients *{τ }* were held fixed at values estimated from the data (assuming *h*^2^ = 0.3 distributed uniformly across baseline annotations) and *χ_s_*^2^ were sampled as 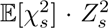 with *Z_s_ ∼ N* (0, 1). We ran four scenarios (null, node-only, edge-only, mixed) with 500 replicates each, *τ* = 5 × 10^−9^ for active annotations and *N* = 340,159. Simulations were conducted independently on heart, brain DLPFC, and gut annotation LD scores to verify calibration across these annotation structures (Supplementary Table 6, sheet “Multi-tissue Simulation”).

#### Power simulation

To map the detection landscape, we conducted a factorial simulation varying five parameters: edge effect size (*τ*_edge_: 0 to 10^−8^, 8 levels), GWAS sample size (*N*_eff_ : 20k–500k, 6 levels), baseline heritability (*h*^2^: 0.05–0.50, 6 levels), node–edge annotation correlation (*r*: 0–0.9, 6 levels), and tissue source (5 tissues). Two-dimensional surfaces (*τ*_edge_ × *N*_eff_ : 48 conditions; *τ*_edge_ × *h*^2^: 48 conditions) were computed in addition to marginal sweeps. Each condition used 200 replicates with the same generative model described above. Power was defined as the rejection rate at *α* = 0.05 (one-sided). To test spatial resolution sensitivity, heart Visium data (4,247 spots) were randomly subsampled to 250, 500, 1,000, 2,000, and 3,000 spots (5 independent subsamples per level), and the full EdgeMap pipeline—spatial graph, communication scoring, LD score annotation, and S-LDSC—was re-run on each subsample before power assessment.

### 4.20 Data sources

Human heart spatial transcriptomics data were obtained from the 10x Genomics Visium platform (4,247 spots; discovery section) and from Kuppe et al. 2022[25] (4 control Visium sections: P1, P7, P8, P17; replication). The Kuppe processed h5ad files also provided the spot-level deconvolution weights used for heart sender–receiver localization. Human brain DLPFC Visium data were obtained from Maynard et al. 2021[18] (discovery: section 151507, 4,226 spots; replication: 11 additional sections 151508– 151676 from 3 donors). Hippocampus Visium data were obtained from Thompson et al. 2025[19] (GSE264692, section V10B01-085 A1, 4,992 spots). Liver Visium data were obtained from Guilliams et al. 2022[20] (GSE192741; discovery: section JBO001; replication: JBO002–JBO004; gene names were converted to human orthologs by uppercase mapping). Cross-resolution validation used cell-segmented Visium HD liver data from Yakubovsky & Itzkovitz 2025[28] (Zenodo 10.5281/zenodo.17735506; samples M2 and M6 from healthy liver donors; 50,278 and 25,671 segmented cells, respectively). These are sample-level segmented-cell counts in the source dataset; the sender–receiver localization below uses the native 8 *µ*m bin matrix for sample M6 (475,519 bins). Intestine Visium data were obtained from Gupta et al. 2024[53] (GEO: GSE189184; discovery: sample B4, 2,575 spots; replication: samples B8, B10, C5).

GWAS summary statistics were grouped as follows. Cardiovascular traits: systolic and diastolic blood pressure from UK Biobank (*N* = 340,159), atrial fibrillation from Nielsen et al.[54] (*N*_eff_ = 228,221), heart rate from UK Biobank (*N* = 340,162), and coronary artery disease from van der Harst & Verweij[55] (*N*_eff_ = 380,832; independent replication: FinnGen Release 10[27], *N*_eff_ = 166,436, no UK Biobank overlap). Psychiatric and neurological traits: schizophrenia from PGC3[56] (*N*_eff_ = 233,471), major depression from Howard et al.[57] (*N* = 807,553), bipolar disorder from PGC3[58] (*N*_eff_ = 101,962), ADHD from PGC[59], and Alzheimer’s disease from Bellenguez et al.[60] (*N* = 788,989). Metabolic and immune traits: BMI from Yengo et al.[61] (*N* = 681,275), ulcerative colitis and Crohn’s disease from de Lange et al.[62] (UC: *N* = 45,975; CD: *N* = 40,266), LDL cholesterol and triglycerides from UK Biobank (independent replication: LDL from the Global Lipids Genetics Consortium[26], *N* = 173,082, no UK Biobank overlap), T2D from FinnGen r10[27], and height from UK Biobank.

HapMap3 SNPs, the SNP–gene weight matrix and the gsMap quick-mode baseline LD scores were obtained from the gsMap resource repository[6, 63]. The regression baseline used in the analyses contains the gsMap quick-mode genic LD-score indicator (all gene) and genome-wide base LD score (base); we therefore describe comparisons to baseline-LD v2.2/v2.3 annotations as a natural future sensitivity analysis rather than as part of the present model.

### 4.21 Cross-section robustness and cross-dataset replication

To assess reproducibility, we ran the full EdgeMap pipeline independently on 12 brain DLPFC sections (151507–151676, from 3 donors[18]), 5 heart Visium sections (1 from 10x Genomics plus 4 control sections P1, P7, P8, P17 from Kuppe et al.[25]; 5 donors), 4 liver Visium sections (JBO001–JBO004, from 4 donors[20]), and 4 gut Visium sections (B4, B8, B10, C5, from 4 donors). Each section was processed independently through spatial graph construction, LR communication scoring, annotation LD score computation, and S-LDSC regression. Cross-section robustness and cross-dataset replication were quantified by a predefined protocol:

#### Layer A (aggregate)

For each tissue–trait pair, we recorded the edge *z*-score in every section. Because serial sections from the same donor are not independent, the primary cross-section summary first averages section-level *z*-scores within each donor and then combines donor summaries by Stouffer’s method; for tissues with one section per donor, this equals section-level Stouffer. We also tested whether donor-level summaries were consistently positive (one-sided binomial sign test). The fraction of sections individually significant at *P* < 0.05 and a DerSimonian–Laird random-effects meta-analysis[64] of section-level *z*-scores are reported as diagnostic sensitivities, not as independent genetic replication statistics.

#### Layer B (context-level gene-set stability)

For independent-GWAS comparisons, we evaluated sign concordance among top-ranked contexts in the discovery GWAS. For cross-section comparisons, we also computed the Spearman rank correlation of context-annotation *z*-score vectors and sign concordance among the top 20 and top 50 contexts ranked by *z* in the discovery section (one-sided binomial test against 50% null), but these cross-section context-level quantities are reported only as diagnostics because they are reproduced by the scale-invariance audit.

#### Layer C (top-tail overlap)

For the same section comparisons, we asked whether the top-*k* contexts in one section were enriched for top-*k* status in the other (*k* = 20, 50). Enrichment was quantified as observed overlap divided by chance expectation (*k*^2^*/N*_pairs_) and tested via one-sided Fisher’s exact test. As with cross-section context-level correlations, this statistic is interpreted as an annotation-structure diagnostic rather than as communication-measurement replication.

For independent-GWAS replication, the same tissue-derived annotation LD scores were paired with substitute GWAS summary statistics (GLGC 2013 LDL[26] for liver; FinnGen Release 10 CAD[27] and CARDIoGRAM 2015[65] for heart) and the full EdgeMap pipeline was rerun; the same three-layer protocol was applied to cross-GWAS context-annotation *z*-score vectors. Because the context tests are constituent-gene annotations, this comparison evaluates gene-set stability across GWAS rather than LR direction or interaction.

### 4.22 Visium HD processing

Cell-segmented Visium HD liver data[28] (samples M2 and M6) were preprocessed using the same Scanpy pipeline as standard Visium (library-size normalization to 10,000, log1p transformation, 5% expression threshold). The spatial neighbor graph was constructed with *k* = 6 nearest neighbors, identical to the standard Visium pipeline. For aggregate edge testing, each segmented cell in the cell-segmented matrix was treated as one spatial unit. Cell-type labels for the sender–receiver analysis (§4.23) were assigned by maximum expression of canonical marker genes: ALB (hepatocyte), ACTA2 (stellate cell), CD68 (Kupffer cell), KRT19 (cholangiocyte), and PECAM1 (endothelial cell).

### 4.23 Sender–receiver localization

For the highlighted liver–LDL maps, we tested whether ligands and receptors localize to distinct cell types using the Visium HD sample M6 section at native 8 *µ*m bin resolution (475,519 bins). Bins were assigned to cell types (hepatocyte, stellate, Kupffer, cholangiocyte, endothelial) based on maximum marker gene expression. For each pair, the “primary sender” was the cell type with the highest mean ligand expression (excluding cell types with zero expression), and the “primary receiver” was the cell type with the highest mean receptor expression. Contexts were labelled cross-cell-type for descriptive mapping when these two cell types differed. This label does not validate signaling, genetic mediation or the named LR relation.

For heart, we used the published spot-level deconvolution weights in the Kuppe et al. control Visium sections rather than assigning each spot to a single cell type. For a directed spatial edge from sender spot *i* to receiver spot *j*, ligand–receptor pair (*L, R*), sender cell type *c_s_* and receiver cell type *c_r_*, we allocated expression-derived LR activity as *W_ij_L_i_R_j_q_i_*(*c_s_*)*q_j_*(*c_r_*), where *W_ij_* is the Gaussian spatial weight and *q_i_*(*c*) is the deconvolution weight of cell type *c* in spot *i*. We summed this quantity over all directed spatial edges to obtain sender–receiver cell-type axes. The baseline for each axis used the same spatial graph and deconvolution weights but omitted ligand and receptor expression, so enrichment is the expression-weighted fraction divided by the cell-type composition fraction. Vascular receiver summaries pooled endothelial, pericyte and vascular smooth-muscle compartments; endothelial receiver summaries used only the endothelial deconvolution weight. This analysis provides compartment-level localization of expression-derived LR scores in Visium spots, not direct physical contact or signaling between individual cells.

### 4.24 Spatial gradient analysis

For each tissue with a nominal-positive aggregate edge estimate, we tested whether expression-derived score maps for within-combination prioritized LR labels vary non-randomly along the tissue’s primary spatial axis. In brain, the spatial axis was defined by cortical layer annotations (layer guess from Maynard et al.). In liver and gut, the axis was defined as the first principal component of spatial coordinates, approximating the portal–central and crypt–villus axes, respectively. In heart, PC1 of spatial coordinates was used.

For each within-combination prioritized LR label (FDR < 0.10 under empirical null calibration), we computed the Spearman correlation between its per-spot expression-derived LR score and the spatial axis coordinate. The aggregate test compared the observed mean absolute correlation across these pairs to a null distribution obtained by permuting spatial axis labels 10^3^ times. The enrichment ratio is the observed mean divided by the permutation mean.

### 4.25 LR-context profile modules

To identify groups of LR contexts with coordinated trait-loading profiles, we constructed a context trait *z*-score matrix from exploratory context-level conditional testing across trait–tissue combinations with nominal-positive aggregate edge association. Contexts present in fewer than two such combinations were excluded. We applied hierarchical clustering (Ward’s linkage, Euclidean distance) to the standardized *z*-score matrix and selected the number of modules *k* that maximized the silhouette score over *k* 5, 6, 7, 8 (optimal *k* = 8, silhouette = 0.309; scores 0.291–0.309 across the grid). Module–trait loadings were computed as the mean context-level *z*-score within each module for each trait.

### 4.26 LR-context annotation-profile correlation (ERC)

For each pair of traits with nominal-positive aggregate edge association in at least one shared tissue, we computed the Spearman correlation of their context-annotation *z*-score vectors within that tissue. When multiple tissues were available, we averaged the within-tissue correlations. The resulting ERC matrix was compared to published genome-wide genetic correlations (*r_g_*) from LD score regression by computing the Spearman rank correlation across all 17 trait pairs for which both ERC and *r_g_* values were available.

### 4.27 Node–edge complementarity analysis

To quantify complementarity between node and edge heritability signals, we computed per-gene GSS and edge communication scores (CSS) for each tissue using the discovery section. For each tissue–trait context contributing FDR < 0.10 candidate rows under empirical null calibration, we extracted the participating genes (“prioritized constituent genes”) and the top-100 genes ranked by node specificity (“node genes”). We quantified overlap via the Jaccard index and assessed whether prioritized constituent genes were ranked differently in the node distribution using the rank-biserial correlation from the Mann–Whitney *U* test. Genome-wide concordance between GSS and CSS was measured by Spearman correlation across all genes.

### 4.28 Cell-type composition baseline control

To test whether edge heritability is confounded by cell-type composition, we augmented the S-LDSC regression with spatial domain annotations. For each tissue, spots were grouped into *K* domains: in brain DLPFC (*K* = 8) we used the published cortical layer assignments[18]; for the other tissues (*K* = 10) domains were identified from expression, by Leiden clustering of a 15-nearest-neighbour graph built on the first 30 principal components of the 2,000 most highly variable genes, with the resolution selected by bisection to reach the target *K*. For each domain *d*, we computed a gene-level specificity score specificity 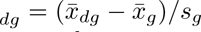, where *x̄_dg_* is the mean expression of gene *g* in domain *d* and *x̄_g_*, *s_g_* are the genome-wide mean and standard deviation. These domain-specificity vectors were mapped to SNP-level LD scores via the same SNP–gene weight matrix **W** used for node and edge annotations, yielding *K* additional annotation columns *ℓ*_domain*,d*_. We then ran S-LDSC with baseline + node + edge + domain annotations and compared edge *z*-scores with and without domain control. Separately, conditioning on gene-level cis-LD burden (a proxy for gene length and local LD architecture) and mean expression—the two most commonly cited technical confounders in LD-score-based enrichment—retained 12 of 13 nominal-positive settings (mean *z*-score change 5.4%), consistent with the domain-conditioning results above.

Because the composition control depends on how domains are defined, we repeated it under three definitions applied uniformly across all five tissues and matched on the number of domains, so that only the definition differs. The first clusters the expression principal components as above. The second uses the raw spot coordinates alone. The third propagates the expression embedding over the spatial six-nearest-neighbour graph before clustering (emb←0.5 emb + 0.5 em̄b*_N_*, two rounds), the step common to SpaGCN[66], STAGATE[67] and BayesSpace[68]; implementing it directly keeps the comparison reproducible without a dependency on any one of those packages. All 13 nominal-positive edge combinations were re-tested under each definition. Eleven of 13 were retained under the expression-derived definition, 12 under the coordinate-only definition and all 13 under the joint definition, with mean edge *z*-score changes of 7.9%, 4.0% and 5.5%, respectively (Supplementary Table 5, sheet “Domain Sensitivity”).

### 4.29 Ligand–receptor specificity test

To test whether the edge annotation captures signal specific to known ligand–receptor biology rather than generic spatial co-expression, we constructed null annotations by replacing each gene in the LIANA Consensus database with a non-LR gene matched on expression level and spatial coefficient of variation. For each tissue, we computed per-gene mean expression *x̄_g_* and spatial CV cv*_g_* = sd(*x_g_*)/(*x̄_g_* + *ɛ*), then partitioned genes into 20 × 20 joint-quantile bins on these two axes. Each LR gene was replaced by a random non-LR gene drawn from the same bin, preserving expression magnitude and spatial variability while removing LR identity. Each shuffled database was processed through the full EdgeMap pipeline (spatial communication, CSS, annotation LD scores, S-LDSC) with the same GWAS and spatial graph as the real analysis. We performed 500 shuffles for each of four trait–tissue combinations (heart–SBP, brain– bipolar disorder, liver–LDL, gut–ulcerative colitis) and computed empirical *P* -values as the fraction of shuffled aggregate edge *z*-scores exceeding the real value.

### 4.30 Additional baseline annotations

To test whether the edge signal is explained by broader gene classes that are themselves enriched for heritability, we constructed four further annotations and added them to the regression, each separately and all together: a constant annotation over all genes; a binary indicator for genes detected in the tissue (12,539–17,625 genes); a binary indicator over the full LIANA Consensus resource (4,624 pairs, 1,877 unique genes, of which 900–1,296 are present per tissue); and a binary indicator for the top decile of mean expression (1,254–1,763 genes). The last is not redundant with the node annotation, because GSS scores spatial concentration rather than expression magnitude: a highly expressed but spatially uniform gene receives a low GSS. Each annotation was mapped to SNP level through the same weight matrix **W**.

Of the 13 nominal-positive edge combinations, 13 were retained after adding the all-gene annotation, 13 after the expressed-gene annotation, 12 after the LR-database annotation, and 12 after the highly-expressed-gene annotation. With all four included simultaneously, 12 of 13 were retained and the mean edge *z*-score fell by 6.3%; the LR-database annotation on its own raises it by 5.3%. The exception in every case is gut–LDL, which carried the weakest original *z*-score. Unlike the broad gene-class baselines, the tissue-active LR indicator is constructed from the same gene support as the edge annotation and is therefore expected to absorb part of the edge estimand. We use it only as a highly conditioned sensitivity analysis. Collinearity means the result is not mathematically ordered below the primary estimate.

### 4.31 Pairing permutation

The preceding test removes ligand–receptor gene identity. As a complementary database-sensitivity analysis, we asked whether the aggregate annotation changes when curated pairing labels are permuted while the ligand and receptor gene universes are held fixed. This is a test of annotation construction, not a genetic test of pair identity. For each tissue the receptor column of the active pair list was randomly reassigned, so that every ligand and every receptor remained present and active but no pair retained its original partner. Communication intensities, pair scores, gene-level CSS and annotation LD scores were recomputed from scratch for each permutation and the regression re-run, with 500 replicates per tissue. Empirical *P* -values are the fraction of permuted aggregate edge *z*-scores exceeding the observed value.

At the level of individual LR labels we used a complementary expression-score partner-resampling diagnostic. For each of the ~1,950 active contexts across the five tissues, the ligand was held fixed and the receptor replaced by each of 500 receptors drawn at random from the database, and PairScore was recomputed. The empirical *P* - value for a pair is the fraction of resampled partners achieving a PairScore at least as large as that of the observed pairing. These expression-derived scores were summarized across contexts by a one-sample *t*-test against the null expectation of 0.5, by the fraction reaching *P* < 0.05 within strata of context specificity, and by the ratio of observed to mean resampled PairScore.

### 4.32 Membership-controlled specification

The edge annotation is built from ligand–receptor genes, so its conditional association can reflect both broad LR-gene membership and the spatially derived scores assigned to those genes. We therefore constructed an arrangement-sensitive gene annotation and conditioned on broad membership and marginal expression. This targeted sensitivity does not isolate or identify a relational genetic effect.

For the pair statistic we used a bivariate spatial cross-correlation. Writing *L̃*^(*p*)^ and *R̃*^(*p*)^ for the mean-centred effective ligand and receptor concentrations of pair *p* across cells,

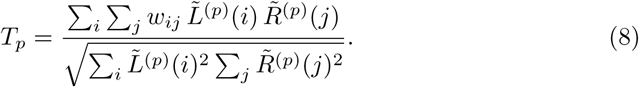

Because both vectors are centred, E[*T_p_*] = 0 whenever the spatial weights *w_ij_* are independent of expression—in particular under permutation of spot coordinates. Pairwise values of *T_p_*, however, have different null variances because ligand and receptor expression profiles differ in sparsity and dynamic range. We therefore standardized each pair against a within-pair permutation null constructed by permuting the ligand vector across spots 500 times while holding the receptor vector and spatial graph fixed (seed 20260625 for the controlled Cross-Moran sensitivity battery):

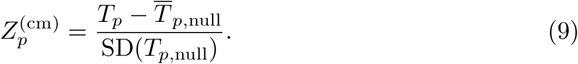

Pairs with zero null variance were assigned *Z_p_*^(cm)^ = 0. The gene-level Cross-Moran score was then defined as the maximum positive standardized pair score incident on that gene, 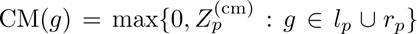. This standardized score responds to spatial alignment alone, whereas PairScore(*p*) measures the concentration of communication intensity relative to its own tissue-wide mean. The two quantities are complementary: concentration identifies which LR contexts are active and spatially localized, whereas cross-correlation forms an arrangement-sensitive gene score for the targeted audit.

Gene-level CM scores were mapped to SNP-level annotations through the same weight matrix **W** used for the node and edge annotations. The controlled regression adds a binary LR-membership indicator and two continuous marginal annotations: natural-scale mean expression for ligand genes and for receptor genes, each mapped through **W** in the same way:

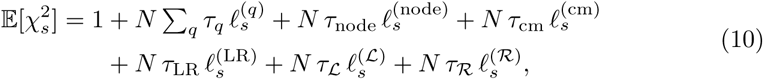

where 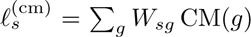 is the Cross-Moran annotation, 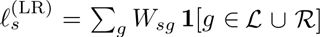 the membership annotation, and 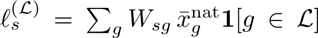 and 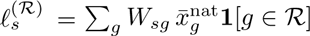 the ligand- and receptor-side marginal annotations. Here 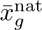 is the mean natural-scale expression of gene *g* across spots.

This computationally intensive sensitivity analysis was implemented on a fixed, non-exhaustive 28-combination case-study and contrast panel: heart with SBP, DBP, CAD, bipolar disorder, LDL, triglycerides and AD; brain with bipolar disorder, schizophrenia, depression, ADHD, AD and height; liver with LDL, triglycerides, T2D, BMI and CAD; gut with UC, Crohn’s disease, CAD, T2D and BMI; and hippocampus with AD, bipolar disorder, schizophrenia, depression and ADHD. The implementation uses this list directly and does not select combinations by their controlled *z*-scores. The panel spans every tissue and the primary case-study and reviewer-motivated positive and negative contrasts, but it is neither a random sample nor an exhaustive analysis of the 85-combination landscape. We therefore interpret it only as a targeted sensitivity analysis and do not extrapolate its retained fraction to the full landscape.

To test whether the residual signal depends on the observed spatial configuration, we permuted spot coordinates, rebuilt the spatial graph and weights *w_ij_*, recomputed *T_p_*, the 500-permutation standardized *Z_p_*^(cm)^, the gene-level CM scores and the annotation LD scores, and re-ran the regression (10 coordinate-shuffle replicates per tissue, seed 20260625). The coordinate permutation is effective at the score level: gene-level CM scores computed from shuffled coordinates correlate with the observed scores at Spearman *ρ* = 0.07–0.18 across tissues.

Across the 28 tissue–trait combinations tested under this specification, two are nominal-positive (liver–TG *z* = 1.89; liver–CAD *z* = 1.70). They are descriptive results within this targeted sensitivity panel, not adjusted discoveries. The coordinate-shuffle diagnostic was evaluated for all 28 combinations. Restricted to the 10 combinations whose frozen controlled *z* exceeds unity—below which a proportional change is uninformative—coordinate shuffling removes a median of 30.5% of the controlled signal, against 2.9% of the signal estimated without the membership terms. The two formal controlled positives show the same direction (liver–TG: 69.2% controlled loss versus 6.2% uncontrolled; liver–CAD: 29.0% controlled loss versus 5.8% uncontrolled), but the full-panel pattern is heterogeneous. We therefore report the coordinate-shuffle result as a diagnostic sensitivity analysis, not as a primary discovery criterion.

We report this specification as a targeted, highly conditioned sensitivity rather than as a lower bound or primary model. The membership, ligand and receptor annotations are built from overlapping gene sets and mapped through the same matrix **W**, and are therefore strongly collinear with the edge annotation. As collinearity predicts, the marginal terms can take negative coefficients and can absorb part of the signal they are intended to partition (for example, the receptor marginal is negative in several heart and brain contrasts; Supplementary Table 6, sheet “Controlled Model”). The specification should therefore be interpreted as a mean-expression marginal control whose coefficient is not guaranteed to be smaller than the primary estimate, not as a binary deletion of ligand and receptor gene sets. Results for all combinations are reported in Supplementary Table 6, sheet “Controlled Model”.

### 4.33 Matched pseudo-pair null test

As a replacement-gene sensitivity analysis, we constructed a matched pseudo-pair null for each top-ranked LR context. This comparison asks whether replacing one constituent with a gene matched on measured properties changes the sparse gene-membership enrichment. It cannot establish pair identity or an LR interaction, because each real and pseudo annotation is still a scaled membership vector. For each pair’s receptor gene, we identified candidate pseudo-receptors from the full gene set, matched on five continuous properties (mean expression, spatial coefficient of variation, cis-LD burden, LR database degree, and GSS) and constrained to the same UniProt sub-cellular localization tier (membrane receptor, secreted ligand, or intracellular). All five properties were standardized to zero mean and unit variance, and candidates within Euclidean distance < 2.0 in this space were retained as the matched pool. For each real pair, 500 pseudo-pairs were generated by sampling from the matched pool (with replacement when pool size < 500), and each pseudo-pair was processed through the same context-level S-LDSC pipeline (baseline + node + pseudo-pair annotation) to obtain a pseudo *z*-score. The matched empirical *P* -value is *P*_matched_ = (*n*_exceed_ + 1)*/*(*N*_pseudo_ + 1), where *n*_exceed_ counts pseudo-pairs whose *z*-score exceeded the real pair’s observed *z*-score. We tested the top 30 context labels per tissue across all available trait–tissue combinations (10 total: heart × SBP/DBP/CAD, brain × SCZ/bipolar, liver × LDL/TG, gut UC, hippocampus SCZ/bipolar). Matching was imperfect: 38 of 50 property-by-combination balance summaries had SMD *>* 0.10 and 16 exceeded 0.50. We therefore treat this only as an exploratory alternative-gene-set sensitivity (Supplementary Table 5, sheets “Alternative Gene Sets” and “Matching Balance”).

### 4.34 Locus ablation test

To test whether edge heritability is driven by a small number of strong GWAS loci, we performed sequential locus clumping: SNPs were sorted by *χ*^2^, the top SNP was designated as a lead, all SNPs within ±500 kb on the same chromosome were assigned to that locus and removed from the pool, and the procedure was repeated to identify *K* independent loci. For each ablation level (*K* 1, 5, 10, 20), we excluded locus SNPs from the GWAS summary statistics and re-ran the S-LDSC regression with the same annotation LD scores. SNP genomic positions were obtained from the HapMap3 LD weight reference files.

### 4.35 Subcellular localization quality control

Ligand–receptor databases curated for cell–cell communication inference occasionally include protein–protein interactions whose partners lack documented extracellular localization (e.g., calmodulin–ion-channel and spectrin–phosphatase complexes that operate intracellularly). Such labels can receive high context-level gene-set statistics when their constituent genes coincide with GWAS loci, even though they are not compatible with extracellular signalling. To distinguish LR labels compatible with extracellular biology from labels lacking such localization, we classified all LIANA Consensus genes by UniProt subcellular location annotations (865 secreted, 928 membrane, 60 intracellular, 24 unknown) and assigned each LR pair to one of three tiers:

1. **Paracrine (secreted → membrane):** ligand classified as secreted, receptor as membrane-anchored. These are localization-compatible curated paracrine labels.
2. **Contact-dependent (membrane → membrane):** both partners membrane-anchored (e.g., neurexin–neuroligin, ephrin–Eph). These are curated contact-dependent labels.
3. **Localization-unverified:** at least one partner is classified as intracellular or unknown (e.g., CALM1/3–CACNA1C, SPTAN1–PTPRA). These pairs are reported for completeness but excluded from biological interpretation.

A small number of genes carry a membrane annotation without being plasma-membrane proteins, because UniProt lists a membrane among several locations. Five were reassigned to tier 3 on manual review and are named here so the step is reproducible: RPSA and MAGED1 (no transmembrane segment; predominantly cytoplasmic and nuclear), PIGA (rough endoplasmic reticulum), RYR2 (sarcoplasmic reticulum; an intracellular calcium-release channel), and FGF13 (an intracellular FGF homologous factor with no signal peptide, which binds the cytoplasmic face of voltage-gated sodium channels).

Tiers 1 and 2 are collectively termed *extracellular-verified* because both partners are localized to the cell surface or secretory pathway; biological interpretation of context-level results (§2.6–§2.8) is restricted to these tiers. As robustness checks, we reran the full EdgeMap pipeline under two filtering stringencies for all 13 nominal-positive aggregate trait–tissue combinations: (i) tiers 1 and 2 jointly (~90% pair retention), retaining 12 of 13 nominal-positive results; and (ii) tier 1 only (54% pair retention), retaining 7 of 13.

### 4.36 Software and reproducibility

The EdgeMap pipeline is implemented in Python using NumPy, SciPy, Scanpy[48], and scikit-learn. The pipeline processes one trait × one tissue in approximately 23 seconds including context-level model fitting; the 50,000-replicate empirical-null calibration is a separate parallel simulation step.

## Data availability

GWAS summary statistics are publicly available: UK Biobank Neale Lab round 2 (http://www.nealelab.is/uk-biobank; SBP, DBP, heart rate, LDL, TG, height); van der Harst and Verweij[55] (CAD); CARDIoGRAMplusC4D[65] (independent CAD replication); Nielsen et al.[54] (atrial fibrillation); GLGC[26] (independent LDL replication); PGC3[56] (SCZ); PGC[58, 59] (bipolar disorder and ADHD); Howard et al.[57] (major depression); Bellenguez et al.[60] (AD); Yengo et al.[61] (BMI); de Lange et al.[62] (UC, Crohn’s disease); FinnGen Release 10[27] (T2D and independent CAD replication). Spatial transcriptomics datasets are available from GEO and ArrayExpress: heart (Kuppe et al.[25], GSE165838; processed Visium h5ad files and deconvolution weights, https://zenodo.org/records/6578047); brain DLPFC (Maynard et al.[18], http://research.libd.org/spatialLIBD/); hippocampus (Thompson et al.[19], GSE264692); liver (Guilliams et al.[20], GSE192741, the spatial subseries of GSE192742); gut (Gupta et al.[53], GSE189184 and SCP1884); cell-segmented Visium HD liver (Yakubovsky and Itzkovitz 2025[28], Zenodo 10.5281/zenodo.17735506). The LIANA Consensus ligand–receptor database (v0.1.13) is available at https://github.com/saezlab/liana. Baseline LD scores and SNP–gene weight matrices are available from the gsMap resource repository[6].

## Code availability

The installable EdgeMap pipeline and its tests are available at https://github.com/cafferychen777/EdgeMap under an open-source license. The revision uses the genomic-order block-jackknife correction in commit 8ebcd46. A versioned publication archive containing the maintained analysis programs, frozen derived CSV/JSON results, separate visualization programs, and an artifact-level provenance manifest is being prepared for deposition. The repository quickstart is an aggregate-workflow installation and file-schema check on fixed synthetic inputs. It requires the separately installed gsMap resource archive, which the script does not download. The fixed synthetic aggregate edge result does not cross the screening threshold, so the quickstart does not enter the conditional context-level branch or reproduce the manuscript’s 50,000-replicate fixed-annotation empirical calibration.

## Acknowledgments

We thank the participants and investigators of all GWAS consortia whose summary statistics made this work possible. Computations were performed on the Texas A&M Department of Statistics Arseven high-performance computing cluster.

## Author contributions

C.Y., X.Z., and J.C. jointly conceptualized the study. C.Y. developed the methodology, implemented the computational framework, conducted formal analysis, and performed data curation, visualization, and validation. X.Z. and J.C. jointly supervised the research, secured funding, provided resources, and served as corresponding authors. All authors participated in manuscript writing and revision.

---

**Extended Data Fig. 1.**
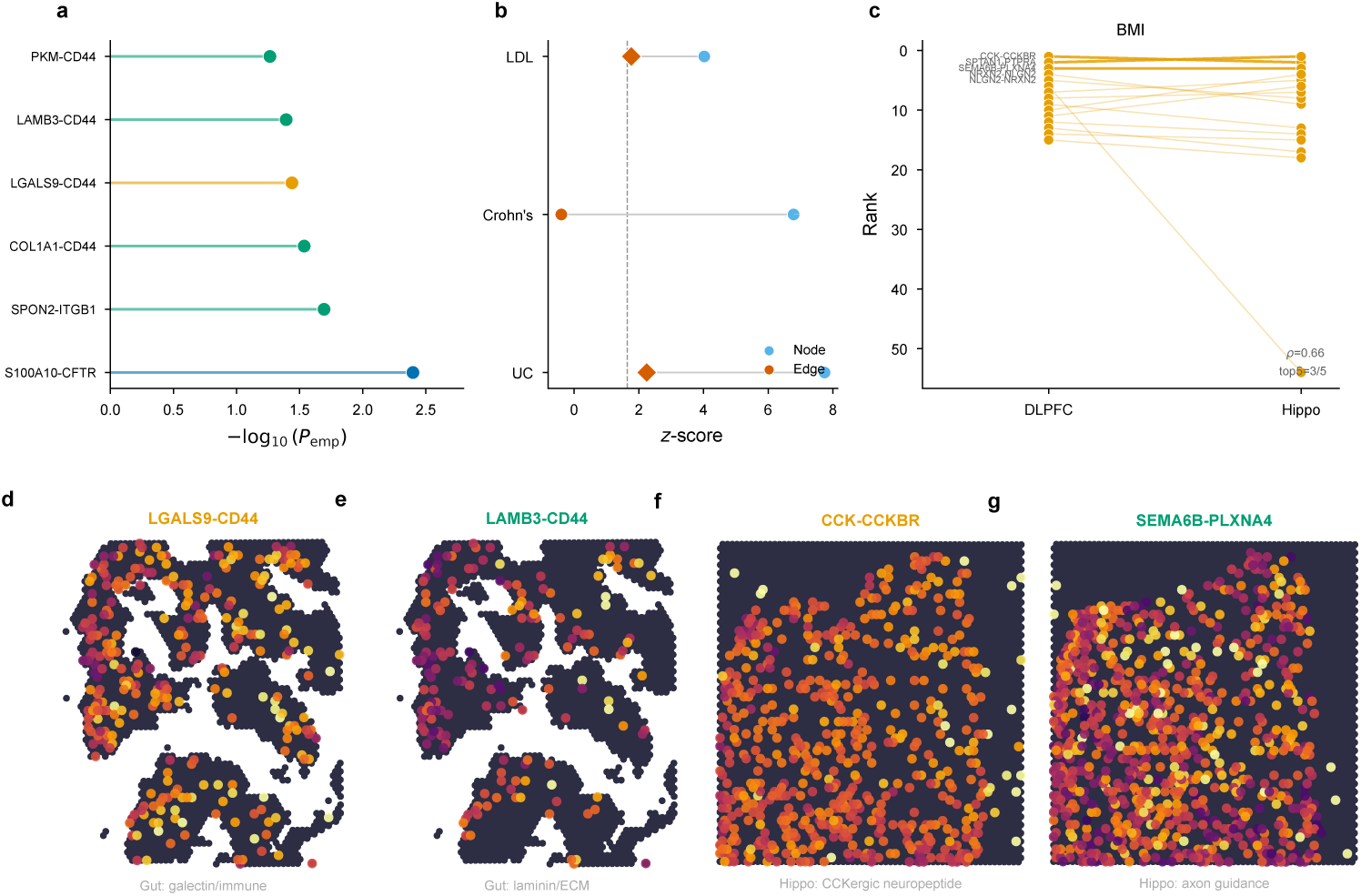
Gut and hippocampus exploratory context summaries. Gut is nominal-positive at the aggregate level for ulcerative colitis but has no within-combination Bonferroni-positive LR-labelled row. **a**, Ranked empirical-*P* summary for gut × UC; labels denote constituent-gene annotations rather than identified LR relations. **b**, Multi-trait comparison in gut. UC and LDL cross the unadjusted one-sided aggregate *P* < 0.05 threshold, whereas Crohn’s disease does not; diamonds denote nominal-positive coefficients. **c**, Descriptive rank concordance among the 15 highest-ranked LR labels between DLPFC and hippocampus for BMI (Spearman *ρ* = 0.66; top-5 overlap = 3/5). **d–e**, Expression-derived spatial LR-score maps for LGALS9–CD44 and LAMB3–CD44 in gut. **f–g**, Corresponding maps for CCK–CCKBR and SEMA6B–PLXNA4 in hippocampus; the maps describe spatial score patterns, not genetic identification of the named relations.

**Extended Data Fig. 2.**
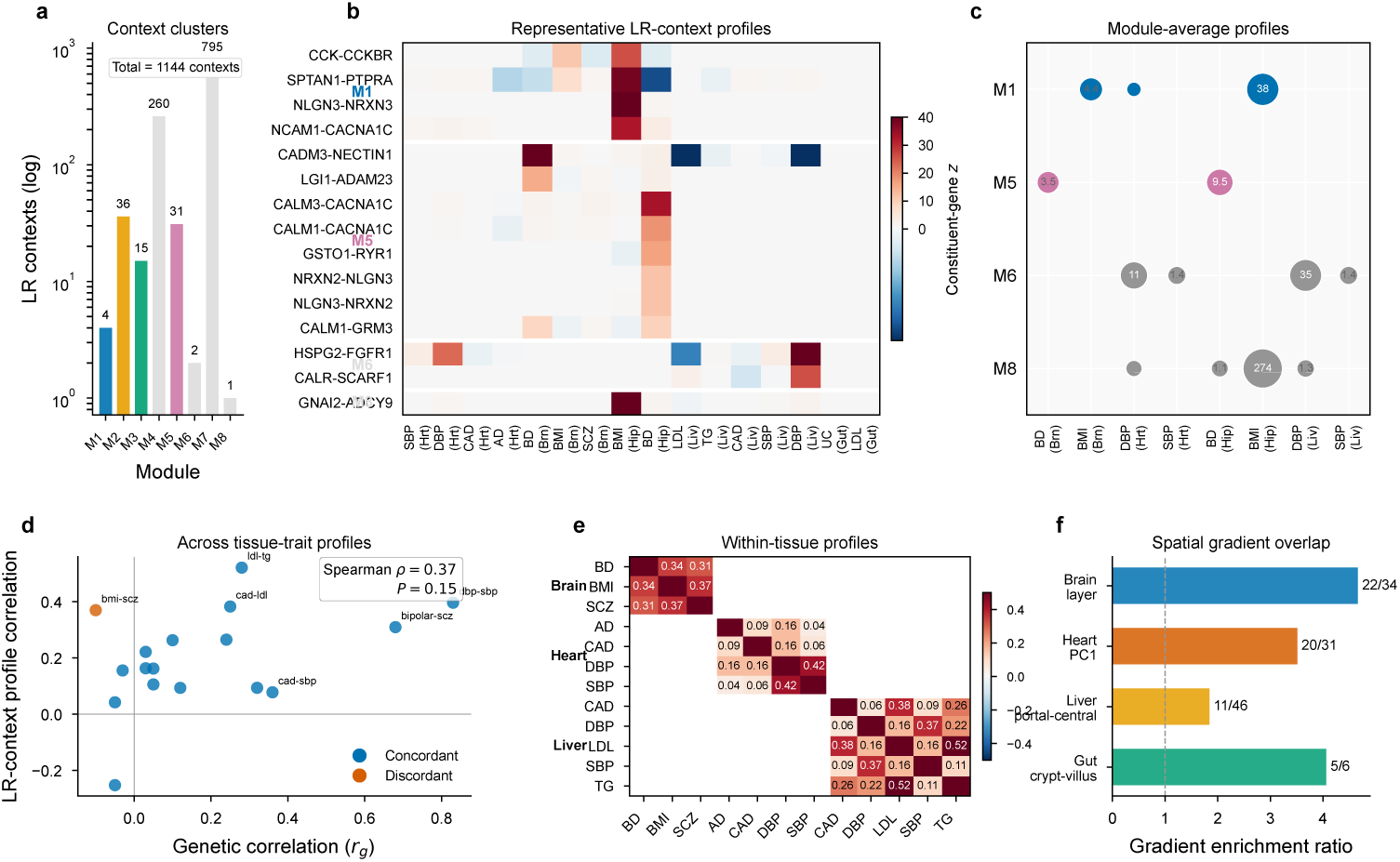
Architecture of LR-context constituent-gene profiles and expression-derived spatial score patterns. **a**, Sizes of the eight frozen context-profile clusters (*n* = 1,144 contexts); the vertical axis is logarithmic. **b**, Constituent-gene annotation *z* profiles for representative contexts from modules M1, M5, M6 and M8 across 16 tissue–trait settings. **c**, Positive module-average *z* profiles for the same four modules. Bubble size increases with the positive module mean, and numbers report the plotted mean. **d**, LR-context profile correlation versus published genome-wide genetic correlation for 17 cross-trait comparisons (Spearman *ρ* = 0.37, *P* = 0.15). Colours distinguish the frozen concordance classification. **e**, Block-diagonal matrices of within-tissue LR-context profile correlations for DLPFC, heart and liver; numbers are off-diagonal correlations. **f**, Overlap of the frozen prioritized-context score maps with tissue spatial gradients. Bars show the observed mean absolute context–gradient correlation divided by its shuffled-axis expectation; labels give the number with nominal rowwise *P* < 0.05 over the number evaluated. Panels **a–e** profile fixed constituent-gene annotations (§2.1).

**Extended Data Fig. 3.**
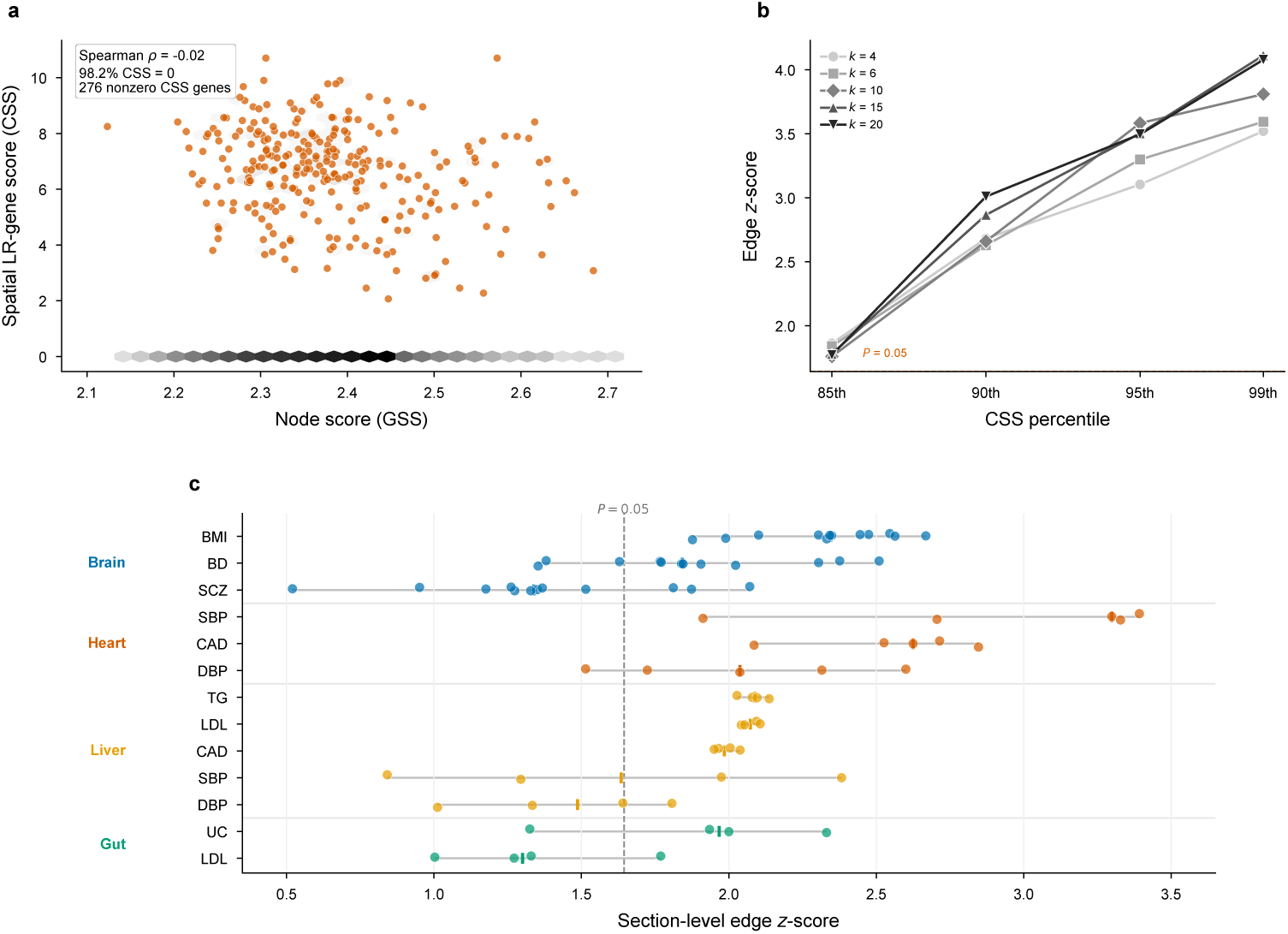
Technical diagnostics: complementarity, parameter sensitivity, and annotation-level stability. **a**, Gene specificity score (GSS, node) vs. context specificity score (CSS, edge) for all 15,217 genes in heart; Spearman *ρ* = *−*0.02, showing low raw score concordance in this tissue. **b**, Parameter sensitivity: edge *z*-scores for heart × SBP across spatial neighborhood sizes (*k ∈ {*4, 6, 10, 15, 20*}*) and context-specificity percentiles (85th–99th). All 20 parameter settings cross the unadjusted one-sided *P* < 0.05 threshold, with *z*-scores increasing monotonically with the percentile threshold. **c**, Cross-section robustness: section-level edge *z*-scores across separate tissue sections (12 DLPFC, 5 heart, 4 liver, 4 gut) for all tested trait–tissue combinations. Points denote sections, horizontal lines denote the observed range, short vertical ticks denote the median, and the dashed line marks *z* = 1.645.

**Extended Data Fig. 4.**
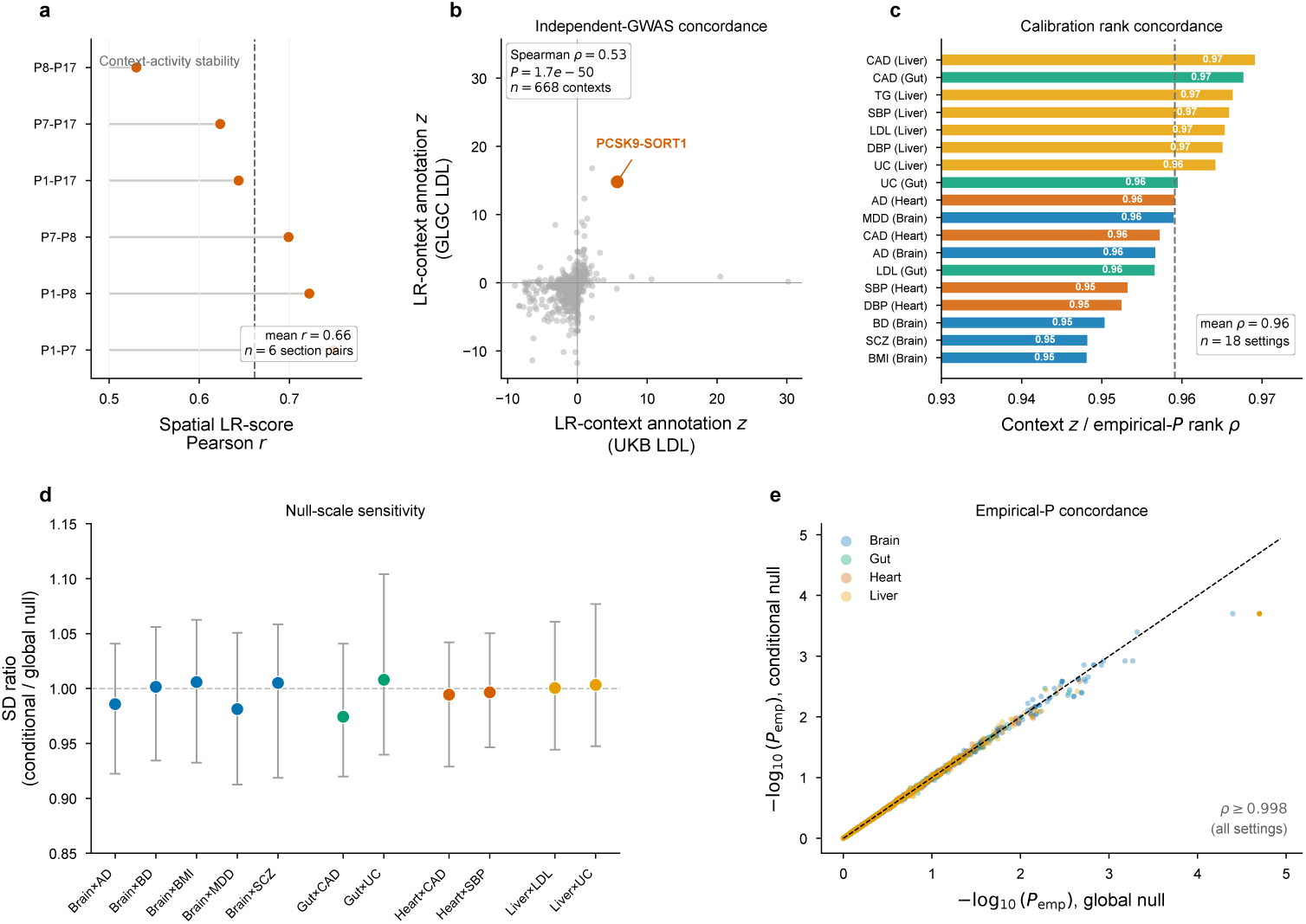
Robustness of fixed LR-context constituent-gene annotation statistics. **a**, Pearson correlation of expression-derived spatial LR scores across the six unique pairs of four Kuppe control-heart sections. Points denote section pairs and the dashed line denotes the mean correlation (*r* = 0.66). **b**, Concordance of 668 liver–LDL constituent-gene annotation *z*statistics between UK Biobank and GLGC 2013 (Spearman *ρ* = 0.53, *P* = 1.7 × 10^−50^); PCSK9–SORT1 is highlighted. **c**, Rank correlation between signed annotation *z* statistics and one-sided empirical-*P* evidence across 18 tissue–trait settings. The mean correlation is 0.96. **d**, Sensitivity of empirical-null scale to conditioning on the observed aggregate cell-intrinsic coefficient. Points are the median conditional/global null standard-deviation ratio for each of 11 settings and bars are interquartile ranges. **e**, Concordance of one-sided empirical *P* values under the global null (*τ*_cell_ = 0) and the conditional null (*τ*_cell_ = *τ̂*_cell_); Spearman *ρ ≥* 0.998 in every setting. Panels **b–e** use fixed constituent-gene annotations (§2.1); these diagnostics assess calibration and reproducibility of constituent-gene statistics.

**Extended data Fig. 5.**
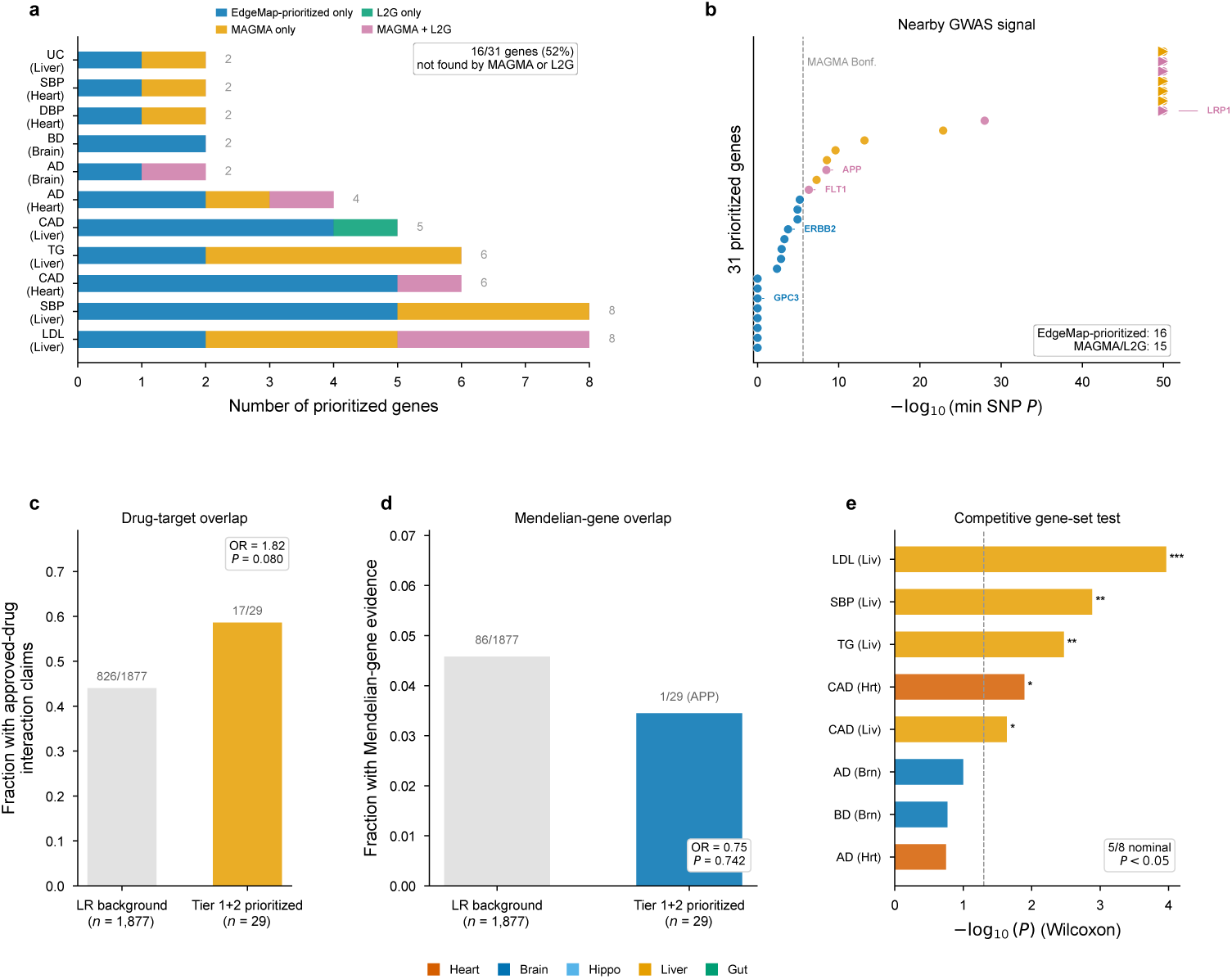
Characterization of prioritized constituent genes. **a**, Counts of prioritized constituent genes by tissue–trait combination and overlap with MAGMA and L2G. Among 31 unique prioritized genes, 16 (52%) were not identified by either comparator. **b**, Strongest nearby GWAS signal for each of the 31 genes, defined as the minimum SNP *P* value within *±*10 kb. Colour denotes MAGMA/L2G overlap status; values above *−* log_10_ *P* = 50 are clipped and shown as triangles. The dashed line marks the MAGMA Bonferroni threshold. **c**, Approved-drug interaction claims among tier 1+2 prioritized genes (17/29) and the LR-gene background (826/1,877); OR= 1.82, unadjusted *P* = 0.080. **d**, Mendelian-gene evidence among tier 1+2 prioritized genes (1/29; APP) and the LR-gene background (86/1,877); OR= 0.75, unadjusted *P* = 0.742. **e**, Competitive Wilcoxon gene-set comparisons between prioritized genes and the LR-gene background in eight testable tissue–trait settings. The dashed line marks unadjusted *P* = 0.05; 5/8 comparisons have nominal *P* < 0.05. Asterisks denote unadjusted *P* < 0.05, *P* < 0.01 and *P* < 0.001. Prioritized genes are derived from fixed constituent-gene annotations (§2.1).

**Extended Data Fig. 6.**
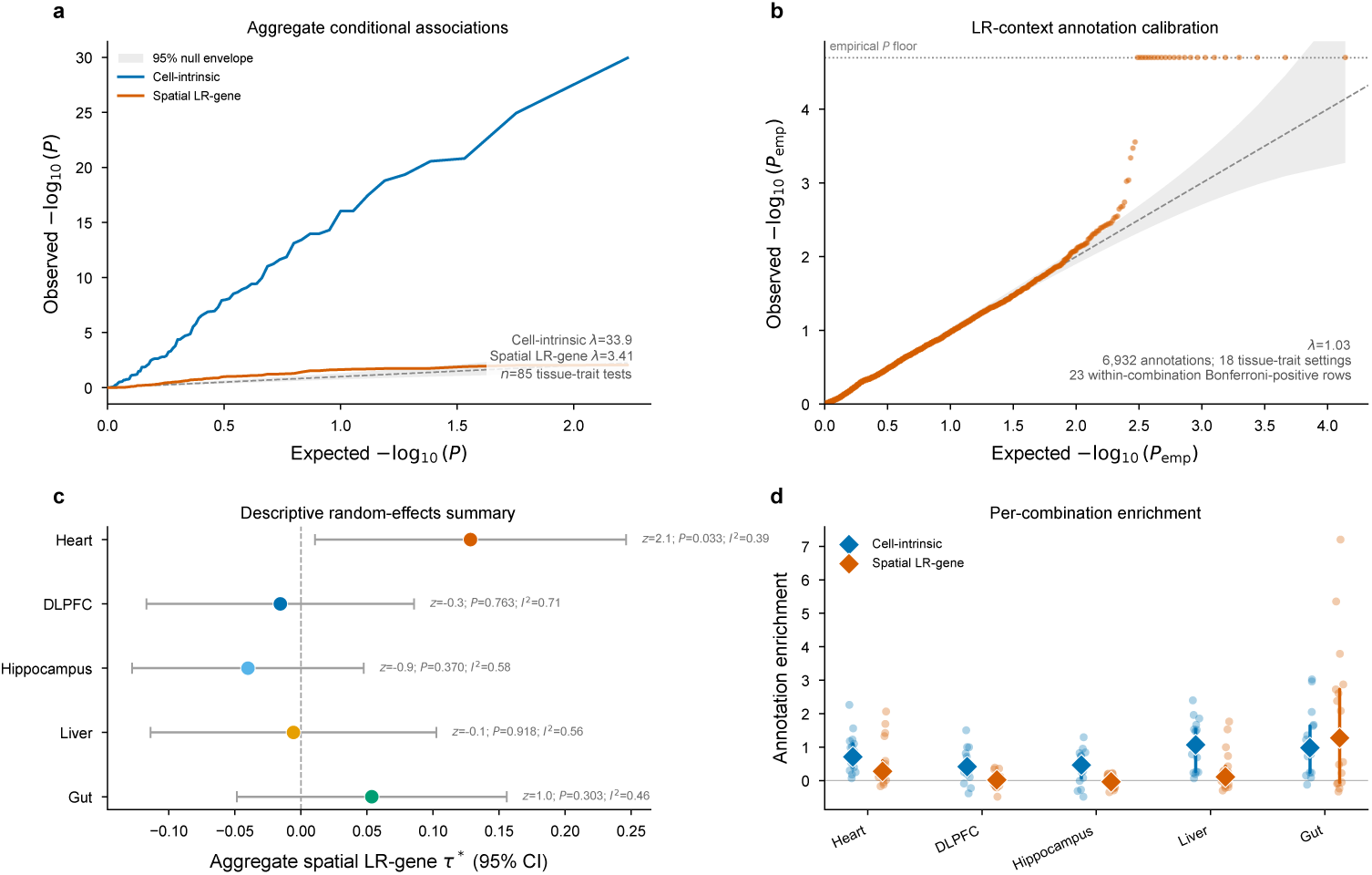
Aggregate effect summaries and LR-context annotation calibration. **a**, QQ plots of unadjusted one-sided aggregate conditional *P* values for cell-intrinsic and spatial LR-gene annotations across 85 tissue–trait tests. The grey region is the pointwise 95% beta null envelope. The displayed *λ* values are descriptive median-*z*^2^ dispersion summaries across the 85 tests, not GWAS genomic-control factors and not evidence of a common effect across tissue–trait combinations. **b**, QQ plot of 6,932 one-sided empirical *P* values for fixed constituent-gene annotations across 18 tissue–trait settings. The horizontal dotted line marks the 50,000-replicate resolution floor, 1*/*(50,000 + 1). Twenty-three rows are within-combination Bonferroni-positive; this count does not apply an additional correction across the 18 settings. **c**, Descriptive random-effects summaries of aggregate spatial LR-gene *τ ^∗^* across 17 traits per tissue. Points are pooled estimates, bars are 95% confidence intervals, and annotations give unadjusted two-sided *P* values and *I*^2^. The heart estimate has *P* = 0.033; the remaining tissue summaries have *P ≥* 0.303. **d**, Cell-intrinsic and spatial LR-gene annotation enrichment for individual tissue–trait combinations. Small points denote combinations, diamonds denote tissue medians, and vertical segments denote interquartile ranges. Panel **b** uses fixed constituent-gene annotations (§2.1); the upper tail ranks these annotations.

**Extended Data Fig. 7.**
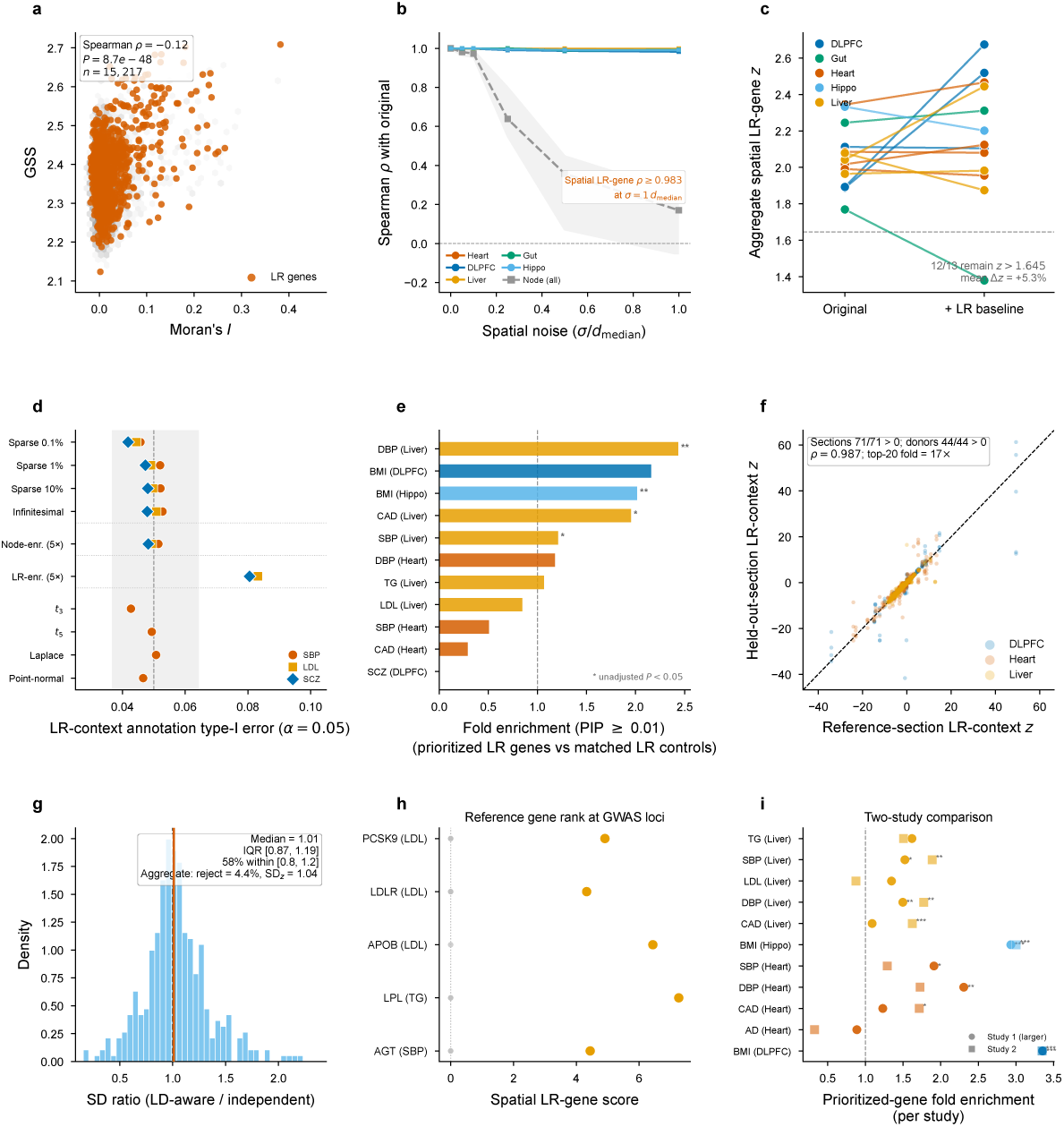
Robustness and complementary characterization of aggregate and constituent-gene estimands. **a**, Moran’s *I* versus gene specificity score for 15,217 heart genes (Spearman *ρ* = *−*0.12, *P* = 8.7 × 10^−48^); LR genes are highlighted. **b**, Robustness to Gaussian coordinate noise across five tissues. Coloured lines show the median correlation of spatial LR-gene scores with the original scores; the dashed grey line and band show the median and interquartile range for cell-intrinsic scores. Spatial LR-gene correlations remain at least 0.983 at one median-neighbour displacement. **c**, Aggregate spatial LR-gene *z* statistics for the 13 nominal-positive combinations before and after adding a binary LR-gene membership baseline. Twelve remain above the nominal one-sided threshold *z* = 1.645; the mean change is +5.3%. **d**, Rejection fractions at *α* = 0.05 for fixed constituent-gene annotations in 22 genotype-derived simulation configurations spanning sparse, infinitesimal, node-enriched, LR-enriched and heavy-tailed architectures. Markers denote SBP, LDL and SCZ where available, the dashed line denotes 0.05, and the grey band is the plotted 95% binomial reference interval. The LR-enriched architecture is a deliberate enrichment stress test, not a pure-null configuration. **e**, Fine-mapped PIP-burden fold enrichment in prioritized LR genes relative to matched LR controls at PIP*≥* 0.01. Eleven of 14 attempted comparisons contain nonzero information; asterisks mark the four comparisons with unadjusted Mann–Whitney *P* < 0.05. **f**, Reference-section versus held-out-section statistics for 5,725 fixed constituent-gene annotation comparisons pooled across DLPFC bipolar disorder, heart CAD, heart SBP and liver LDL. The inset reports summaries from the broader frozen replication protocol: all 71 aggregate section estimates across 11 replicable combinations were positive; at the independent donor–trait level, 44 of 44 were positive (sign-test *P* = 5.7 × 10^−14^). Mean annotation-rank correlation across 41 section comparisons was 0.987, and mean top-20 overlap enrichment was 17×. These are reproducibility summaries for fixed gene-set statistics, not independent replication of LR-pair identities. **g**, Distribution of null standard-deviation ratios for LD-aware versus independent simulations across 370 annotations (2,000 replicates per null scheme). The median is 1.01 (IQR 0.87–1.19). A separate 1,000-replicate aggregate pure-null simulation gives a one-sided rejection fraction of 4.4% and SD(*z*) = 1.04. **h**, Spatial LR-gene scores for genes at five established liver GWAS loci. Orange points identify the reference genes; grey points are other genes in the corresponding locus windo_5_w_5_s. The reference gene ranks first in each displayed locus. **i**, Prioritized-gene PIP-burden fold enrichment in the two largest available studies for each of 11 tissue–trait settings. Circles and squares denote the two studies and asterisks denote unadjusted Mann–Whitney *P* values. Potential cohort overlap precludes interpreting this panel as independent replication. Context-level statistics in panels **d** and **f** use fixed constituent-gene annotations (§2.1).

**Extended Data Fig. 8.**
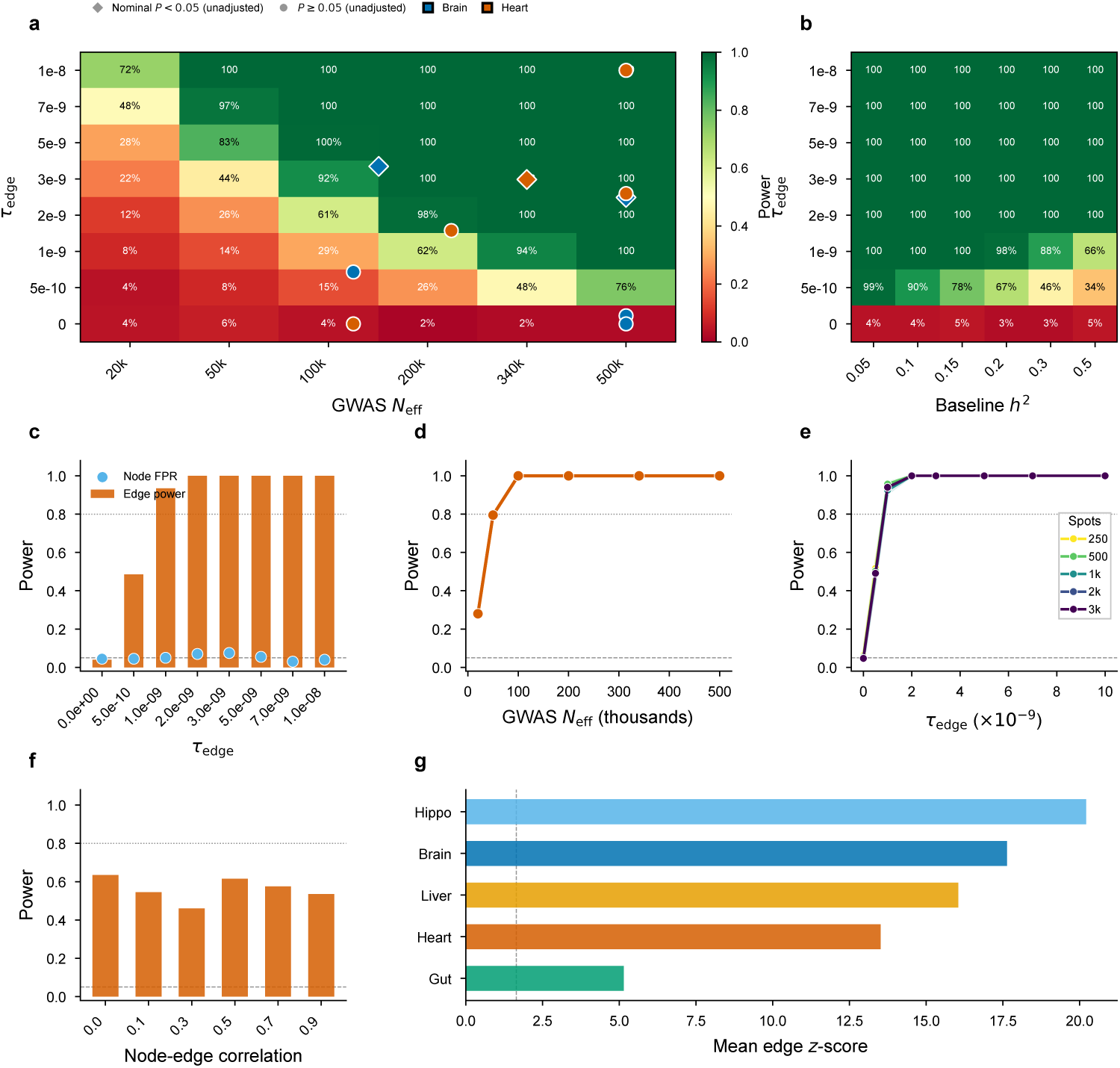
Power and failure-mode simulation. **a**, Power (rejection rate at *α* = 0.05; 200 replicates per cell) as a function of true *τ*_edge_ and GWAS *N*_eff_ (*h*^2^ = 0.3). Diamonds and circles: 10 real tissue–trait results with rebuilt estimates (3 nominal-positive, 7 below the unadjusted threshold), coloured by tissue. Points outside the plotted *N*_eff_ or *τ*_edge_ range are clipped to the panel boundary. **b**, Power as a function of *τ*_edge_ and baseline heritability (*N*_eff_ = 340k). **c**, Marginal *τ*_edge_ sweep (*h*^2^ = 0.3, *N*_eff_ = 340k): edge power (bars) and node false-positive rate (points) at *α* = 0.05. **d**, *N*_eff_ power curve at *τ*_edge_ = 5 × 10^−9^. **e**, Real tissue subsampling: heart Visium subsampled to 250–3,000 spots (5 subsamples each, full pipeline re-run per subsample). Power curves at different spot counts show minimal practical differences; null false-positive rate 4.5–6.2%. **f**, Power as a function of node–edge annotation correlation (*r* = 0.0–0.9). **g**, Tissue comparison: mean edge *z*-score under edge-only enrichment (*τ* = 5 × 10^−9^) across five tissues.

